# Progressive changes in hippocampal stem cell properties ensure lifelong neurogenesis

**DOI:** 10.1101/2020.03.12.987107

**Authors:** Lachlan Harris, Piero Rigo, Thomas Stiehl, Zachary Gaber, Sophie H. L. Austin, Maria del Mar Masdeu, Amelia Edwards, Noelia Urbán, Anna Marciniak-Czochra, François Guillemot

## Abstract

Neural stem cell numbers fall rapidly in the hippocampus of juvenile mice but stabilise during adulthood, ensuring lifelong hippocampal neurogenesis. We show that this reduction in stem cell depletion rate is the result of multiple coordinated changes in stem cell behaviour. In particular, while active neural stem cells divide only once or twice before differentiating rapidly in juveniles, they increasingly return to a resting state of shallow quiescence and progress through additional self-renewing divisions in adulthood. Single-cell transcriptomic, mathematical modelling and label-retention analyses indicate that resting cells have a higher activation rate and greater contribution to neurogenesis than dormant cells, which have not left quiescence. These progressive changes in stem cell behaviour result from reduced expression of the pro-activation protein ASCL1 due to increased post-translational degradation. These mechanisms help reconcile current contradictory models of hippocampal NSC dynamics and may contribute to the different rates of decline of hippocampal neurogenesis in mammalian species including humans.

## INTRODUCTION

Neural stem cells (NSCs) are found throughout the brain during embryogenesis but persist only in restricted brain areas during adulthood, mostly in the dentate gyrus (DG) of the hippocampus and the ventricular-subventricular zone (V-SVZ) of the lateral ventricles, where they produce new neurons that contribute to the functions of the hippocampus and olfactory bulb, respectively. In mice, proliferative dentate precursor cells enter quiescence during the 2^nd^ week after birth. By the end of the second postnatal week, quiescent dentate precursors have acquired the elongated and branched morphology of adult NSCs and the DG has acquired its adult structure (Li et al., 2013; Nicola et al., 2015; Noguchi et al., 2019; Berg et al., 2019). There is therefore a clear transition from developmental neurogenesis, that extends from mid-embryogenesis to the second postnatal week, to adult hippocampal neurogenesis that starts around postnatal day 14 (P14) and continues throughout life (Hochgerner et al., 2018; Noguchi et al., 2019; Berg et al., 2019).

However, hippocampal neurogenesis continues to change considerably after P14 in mice. Indeed, a stereotypic pattern has been observed in most mammals, whereby high numbers of new DG neurons are generated in juveniles, their numbers then decline rapidly in young adults and are then maintained at low levels after sexual maturity and throughout adulthood (Amrein et al., 2015; Snyder, 2019). This transition from high levels of hippocampal neurogenesis in juveniles to lower and sustained levels in adults differs from an aging process as it occurs early in life (Encinas et al., 2011), suggesting that it is adaptive. Interestingly, whether the same transition occurs in humans to allow neurogenesis to extend beyond childhood has recently been disputed, highlighting the lack of understanding of the processes that preserve neurogenesis between infancy and old age (Boldrini et al., 2018; Kempermann et al., 2018; Moreno-Jimenez et al., 2019; Paredes et al., 2018; Sorrells et al., 2018; Spalding et al., 2013).

Hippocampal NSCs sit atop the lineage hierarchy of neurogenesis and changes in their behaviour are likely involved in the transition towards lower and sustainable levels of neurogenesis during adulthood. However, the behaviour of adult hippocampal NSCs is still not well understood and current divergent models are not yet reconciled (Lazutkin et al., 2019). For example, Encinas et al. (2011) described a ‘disposable stem cell model’ whereby hippocampal NSCs that have left quiescence progress rapidly through a series of neurogenic divisions without returning to quiescence and eventually differentiate into astrocytes, leading to a decrease in the NSC pool. Bonaguidi and colleagues (2011) proposed a contrasting ‘long-term self-renewal’ model whereby many hippocampal NSCs can return to quiescence after dividing so that the NSC pool declines only slightly with age. Finally, we recently proposed a model supporting heterogeneity in stem cell behaviour, whereby the degradation of the pro-activation factor ASCL1 in proliferating stem cells allowed a subset of these cells to return to quiescence (Urban et al., 2016).

Here, we show that these different models are all valid but describe NSC behaviour at different stages throughout adult development. Hippocampal NSCs undergo dramatic changes to their properties in early adulthood that serve to preserve the NSC pool. At the onset of adult neurogenesis, all proliferating NSCs are rapidly lost through differentiation, while by six months of age more than half of dividing NSCs return to quiescence. Through mathematical modelling, single-cell RNA sequencing and label-dilution analysis, we determine that cells that have returned to quiescence (resting NSCs) have distinct molecular and cellular properties from quiescent cells that have never proliferated (dormant NSCs) and play an increasingly important part in maintaining NSC proliferation over time. Finally, we find that these coordinated changes are caused by the increased post-translational degradation of ASCL1 protein. Our results demonstrate that progressive and coordinated changes in NSC properties enable a transition from high levels of stem cell proliferation coupled with exhaustion towards lower but sustainable levels of proliferation throughout adult life.

## RESULTS

### Rapid NSC depletion in juveniles and reduced depletion in adults

We set out to examine the cellular properties of hippocampal NSCs throughout adulthood. We began by counting NSCs in the DG of C57Bl/6 mice from the end of perinatal development to old age. We identified NSCs as cells containing a radial GFAP-positive process linked to a SOX2-positive nucleus in the subgranular zone (SGZ). These cells were defined as quiescent or active depending on expression of the cell-cycle marker Ki67. At the onset of adult neurogenesis in 0.5-month old mice (Berg et al., 2019; Noguchi et al., 2019), there were approximately 46,000 NSCs per DG, which underwent an immediate and rapid decline so that their number had more than halved to 17,000 NSCs/DG in 2-month old mice (Figure 1A, B). After this rapid loss of NSCs in juvenile mice, the depletion rate defined as the percentage of the hippocampal NSC pool that was lost each day, slowed dramatically in young adults and depletion continued at a much lower rate in 6-, 12- and 18-month-old mice. The disposable stem cell model proposed for hippocampal NSCs (Encinas et al., 2011) states that NSC activation leads to a series of asymmetric divisions and the eventual loss of the NSC through astrocytic (Encinas et al., 2011) or neuronal (Pilz et al., 2018) differentiation. Since this model links the rate of depletion to NSC activity, a reduced activity should result in a lower rate of NSC depletion. Indeed, the depletion rate decreased in proportion to the size of the active NSC pool in juveniles (between 1 and 2 months, Figure 1C, D). However, the depletion rate slowed down considerably more in young adults (between 2 and 6 months of age) and became decoupled from proliferation, so that it became substantially lower than predicted by the disposable stem cell model (Figure 1D) (Lazutkin et al., 2019). We therefore reasoned that active NSCs may acquire distinct properties in early adulthood that function to slow down depletion and preserve the NSC population.

**Figure 1:**
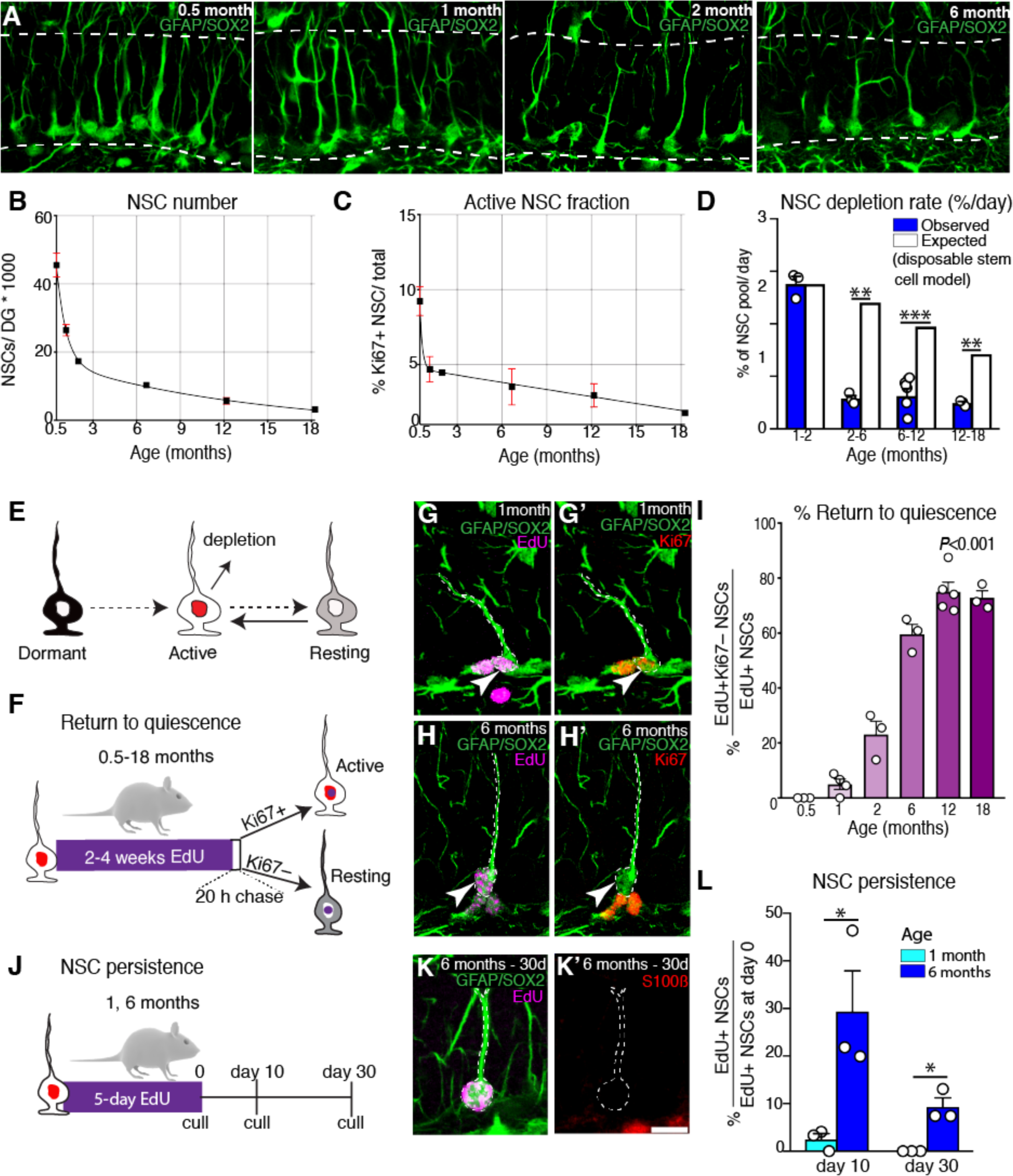
Active NSCs increasingly return to quiescence with time. (**A**) Images and (**B**) quantification demonstrating that the loss of hippocampal NSCs is rapid in young mice (0.5-2 months) but more gradual in adults (> 2 months of age). **(C)** The proliferative activity of NSCs declines slowly with age but the rate of NSC loss (**D**) is lower than predicted by the disposable stem cell model after 2 months of age. (**E**) Schematic of the different hippocampal NSC states: dormant NSCs have never activated while resting NSCs have returned to quiescence from an actively proliferating state. (**F**) Active NSCs were labelled in young and older animals either via EdU-injections (at 0.5 months of age) or through EdU-drinking water for 2-weeks (at 1, 2 and 6 months of age) or 4-weeks (at 12-18 months of age) followed by a 20 h chase. (**G**, **I**) In young mice, EdU+ NSCs rarely returned to quiescence (Ki67-), (**H**, **I**) while return to quiescence increased in frequency with age. The length of EdU administration had a negligible effect on the measured rate of return to quiescence (data not shown). (**J**) The ability of EdU-incorporating NSCs in 1- and 6-month-old mice to persist in the hippocampal niche was determined by measuring the fraction of EdU+ NSCs that remained as NSCs 10 and 30-days post a 5-day EdU labelling protocol. (**K**, **L**) EdU+ NSCs persisted for longer in 6-month-old mice than 1-month-old mice. Graphs show mean ± s.e.m. Dots represent individual mice. Statistics: Student’s t-test in **D**, **L** and one-way ANOVA in **I**. \**P*<0.05, ** *P* < 0.01, \*\*\**P*<0.001. Scale bar (in **J’**): A = 19 μm, **F**, **G** = 14.2 μm, **J**-**J’** = 11.25 μm.

### Active NSCs progressively acquire the capacity to return to quiescence

A mechanism by which active hippocampal NSCs might avoid depletion in young adults could be by returning to quiescence. To address this possibility, we examined the frequency with which activated hippocampal NSCs return to a quiescence state throughout adulthood. We refer to active NSCs that have returned to quiescence as resting NSCs, to distinguish them from dormant NSCs that have remained quiescent since establishment of the DG niche during the first two postnatal weeks (Figure 1E) (Urban et al., 2016). To label active hippocampal NSCs, mice were given the thymidine analogue EdU, either through a single intraperitoneal injection to 2-week-old animals or for periods of 2-4 weeks in their water for older mice (Figure 1F, I) (Chehrehasa et al., 2009). Mice were then culled after a 20-h chase-period and EdU-positive NSCs were identified as having remained cycling during the chase period or having returned to a quiescent state depending on their co-expression of Ki67 (Fig 1F). In 2-weeks old mice, the hippocampal NSC niche is fully mature and comprises a large population of mostly quiescent NSCs (∼90%, Figure 1C) (Berg et al., 2019; Noguchi et al., 2019). However all EdU+ NSCs in 2-week-old mice remained cycling at the end of the chase period (EdU+ Ki67+; 134/134), indicating that at the onset of adult neurogenesis, active NSCs do not have the capacity to return to quiescence and thus adhere closely to the disposable stem cell model (Figure 1G, I).

In 1-month-old mice in contrast, we found that a small fraction (5.01±1.88%) of EdU+ NSCs returned to quiescence (EdU+ Ki67-; Figure 1H, I). The fraction of active NSCs that exited the cell cycle then increased in a gradual and substantial manner so that the majority of EdU+ NSCs returned to a quiescent state in mice at 6-months of age and older (Figure 1I). We confirmed these results with an independent approach to label active NSCs utilising mice containing a *Ki67-creER^T2^* allele (Basak et al., 2018) and a *lox-STOP-lox-tdTomato* cre-reporter (Madisen et al., 2010), hereafter referred to as Ki67^TD^ mice. Transcription of *Mki67* and expression of creER^T2^ occurs in cells progressing through the cell cycle and is excluded from quiescent NSCs (Miller et al., 2018). We administered tamoxifen to 1- and 6-month-old Ki67^TD^ mice for 5 days to label active NSCs and their progeny and culled the mice at the end of the injection period. Consistent with the EdU-labelling experiment, active NSCs (tdTomato+ GFAP+) returned to a quiescent state (Ki67-) at a greater rate in 6-month-old mice than in 1-month-old Ki67^TD^ mice (Figure S1).

We reasoned that because activated NSCs increasingly return to a quiescent state rather than differentiating, they might persist as NSCs for longer in older mice, helping to preserve the NSC pool. To examine the persistence of active NSCs in the hippocampal niche, we labelled active NSCs with EdU for 5-days in drinking water and examined their fate after 10- and 30-days (Figure 1J). We determined that an EdU+ cell had retained its NSC identity if it displayed a radial GFAP+ process and a SOX2+ nucleus in the SGZ and was negative for the astrocytic marker S100B (Martin-Suarez et al., 2019). In 1-month-old mice, only 2.46±1.25% of EdU+ NSCs retained their NSC identity 10-days after labelling and none retained it after 30-days (Figure 1L). Strikingly, in 6-month-old mice, there was an approximate 15-fold increase in the fraction of proliferating NSCs that persisted as NSCs after the 10-day chase (29.37±8.5%) and a third of those (9.3±1.89%) retained their NSC identity after the 30-day chase (Figure 1K, L). These findings demonstrate that active hippocampal NSCs progressively acquire the capacity to return to quiescence, resulting in NSCs persisting in the niche rather than depleting and supporting the maintenance of the NSC pool with age. Importantly, we found that these changes in NSC behaviour arise early (in mice as young as 1-month of age; Figure 1I) and take place mostly during the first 12 months of life. These changes are therefore distinct from ageing-associated pathological changes that lead to NSC dysfunction (Ibrayeva et al., 2019; Leeman et al., 2018; Miranda et al., 2012; Seib et al., 2013; Yousef et al., 2015)

### Resting NSCs increasingly contribute to the proliferative NSC pool

The finding that active hippocampal NSCs increasingly return to a resting state raised the possibility that the dormant population of NSCs, which have not previously proliferated, may also alter their properties with time. To address this, we re-analysed the mice that received prolonged EdU exposure in the drinking water for 2-4 weeks and that were culled after a 20-hour chase (Figure 2A). Resting NSCs do not persist long-term, particularly in young mice (Figure 1L) and therefore most active and resting NSCs are labelled by prolonged EdU exposure in these paradigms, meaning that the EdU-negative NSC population essentially corresponds to the dormant NSC pool. We found that the fraction of dormant NSCs that became active during the 20-hour chase period (i.e. EdU-Ki67+ NSCs) declined progressively and substantially with age, indicating that dormant NSCs were progressing into a deeper state of quiescence (Figure 2B, C). This observation is consistent with the analysis of quiescence in fibroblast cultures (Kwon et al., 2017) and with recent modelling predictions and experimental data showing that NSCs in the adult ventricular-subventricular zone (V-SVZ) deepen their quiescence with age (Bast et al., 2018; Kalamakis et al., 2019; Ziebell et al., 2018).

**Figure 2:**
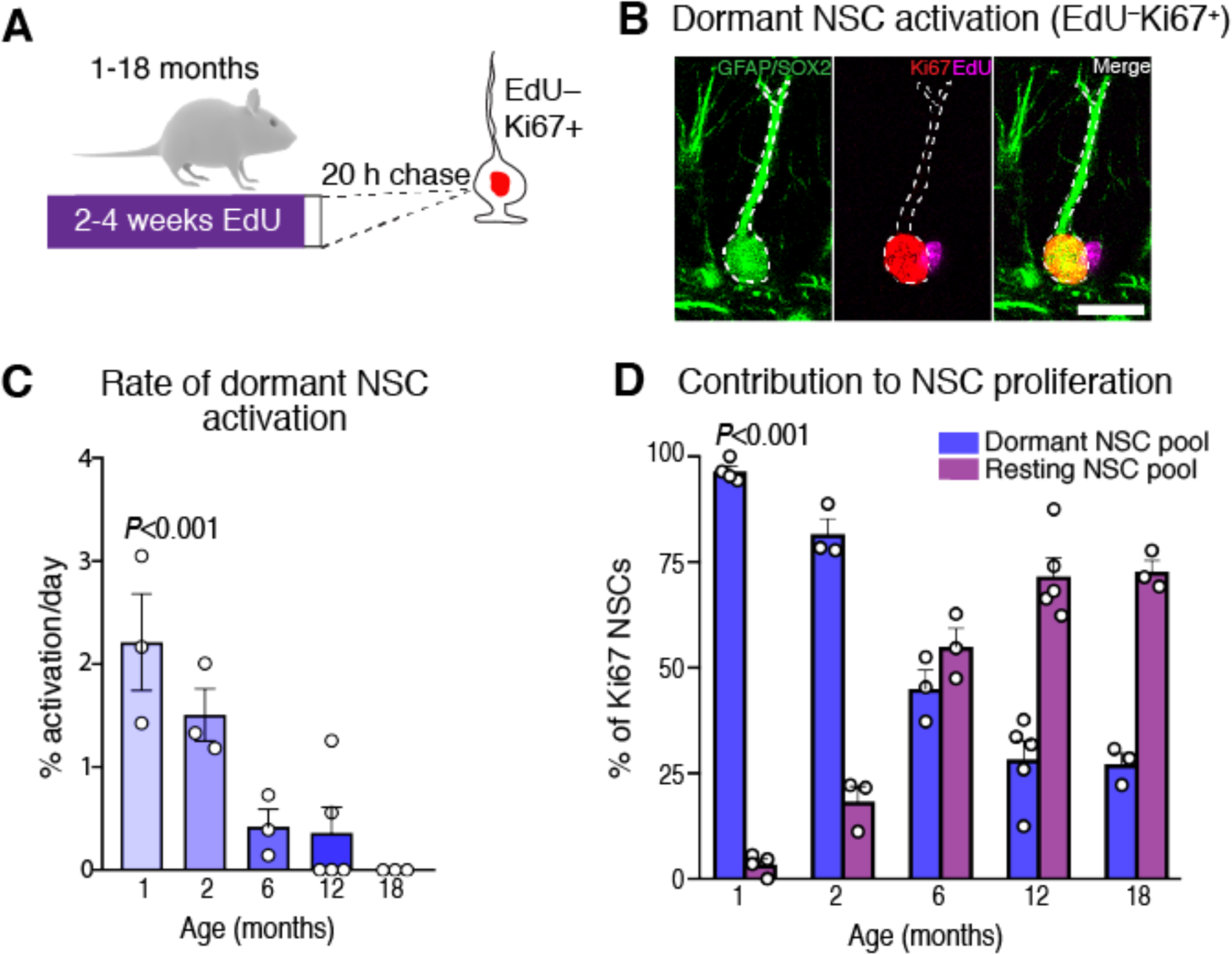
Resting NSCs increasingly contribute to the proliferative NSC pool with time. (**A**) Mice received EdU via drinking water for either 2-weeks (at 1, 2 and 6 months) or 4-weeks (at 12, 18 months) to label active and resting NSCs, followed by a 20 h chase. The length of the EdU administration was increased with age to account for the increased persistence of active and resting NSCs in old mice. (**B**) Dormant NSCs that activated during the 20 h chase period were identified as EdU-Ki67+ NSCs. (**C**) The activation rate of dormant NSCs decreased with age indicating the deepening quiescence of this pool. (**D**) The contribution of dormant NSCs to the proliferative NSC pool decreased with age while the contribution of resting NSCs increased. Graphs show mean ± s.e.m. Dots represent individual mice. Statistics: One-way ANOVA in **C**, Two-way ANOVA in **D**. Scale bar (in B): 10 μm.

However even in 6-12-month-old mice, the numbers of resting NSCs remained small compared to the size of the dormant pool. We therefore assessed the functional significance of resting NSCs by determining their contribution to the maintenance of the proliferative NSC pool at different ages (see Materials and Methods). As expected, in young mice where active NSCs rarely return to quiescence, the vast majority of proliferating NSCs arose from the dormant NSC pool (Figure 2D). However, in 6-18-month-old mice, resting NSCs were the origin of 55-73% of proliferating NSCs (Figure 2D), despite comprising only 3-5% of the NSC population. These findings support a novel model whereby, with time, hippocampal NSC proliferation is increasingly generated from resting NSCs that have the capacity to shuttle between active and quiescent states, thereby delaying NSC depletion and ensuring the long-term maintenance of the NSC pool. These findings also imply that resting hippocampal NSCs have a higher rate of activation than dormant NSCs.

### Increasing numbers of NSC self-renewing divisions over time

Thus far we have demonstrated that the disposable stem cell model (Encinas et al., 2011) describes hippocampal NSC behaviour in juvenile mice but that it no longer wholly applies when mice reach adult ages. Instead, NSCs increasingly approach the behaviour described by Bonaguidi and colleagues (2011) where cells return to quiescence and maintain their identity over the longer-term (Figure 1D, I, L). We next wondered whether returning to the resting state and escaping differentiation might provide NSCs with additional opportunities for self-renewing divisions over time. To address this possibility, we developed a mouse model to count the number of self-renewing divisions that each hippocampal NSC undergoes prior to depleting from the niche. We used a labelling system based on the Histone 2B (H2B)-Green Fluorescent Protein (GFP) fusion protein, which accumulates in nuclei and is then repressed through doxycycline (Dox) administration, resulting in dilution of the GFP label following each cell division (Kanda et al., 1998; Tumbar et al., 2004). We compared juvenile (1.5 month) and adult (6 month) cohorts, in which the nuclear label was targeted to NSCs using the *Glast* gene promoter (Figure 3A-F) (Mori et al., 2006). Most NSCs underwent between 0 and 3 self-renewing divisions before depleting and losing their stem cell identity (Figure 3G, H). Importantly the majority of NSCs in juvenile mice self-renewed only once (59% versus 47% in adults), while most NSCs in adult mice underwent two or more self-renewing divisions (52% *versus* 38% in juveniles) (Figure 3H). The increased self-renewal of hippocampal NSCs with time represents an additional mechanism contributing to the preservation of the NSC pool beyond the juvenile period and throughout adulthood.

**Figure 3:**
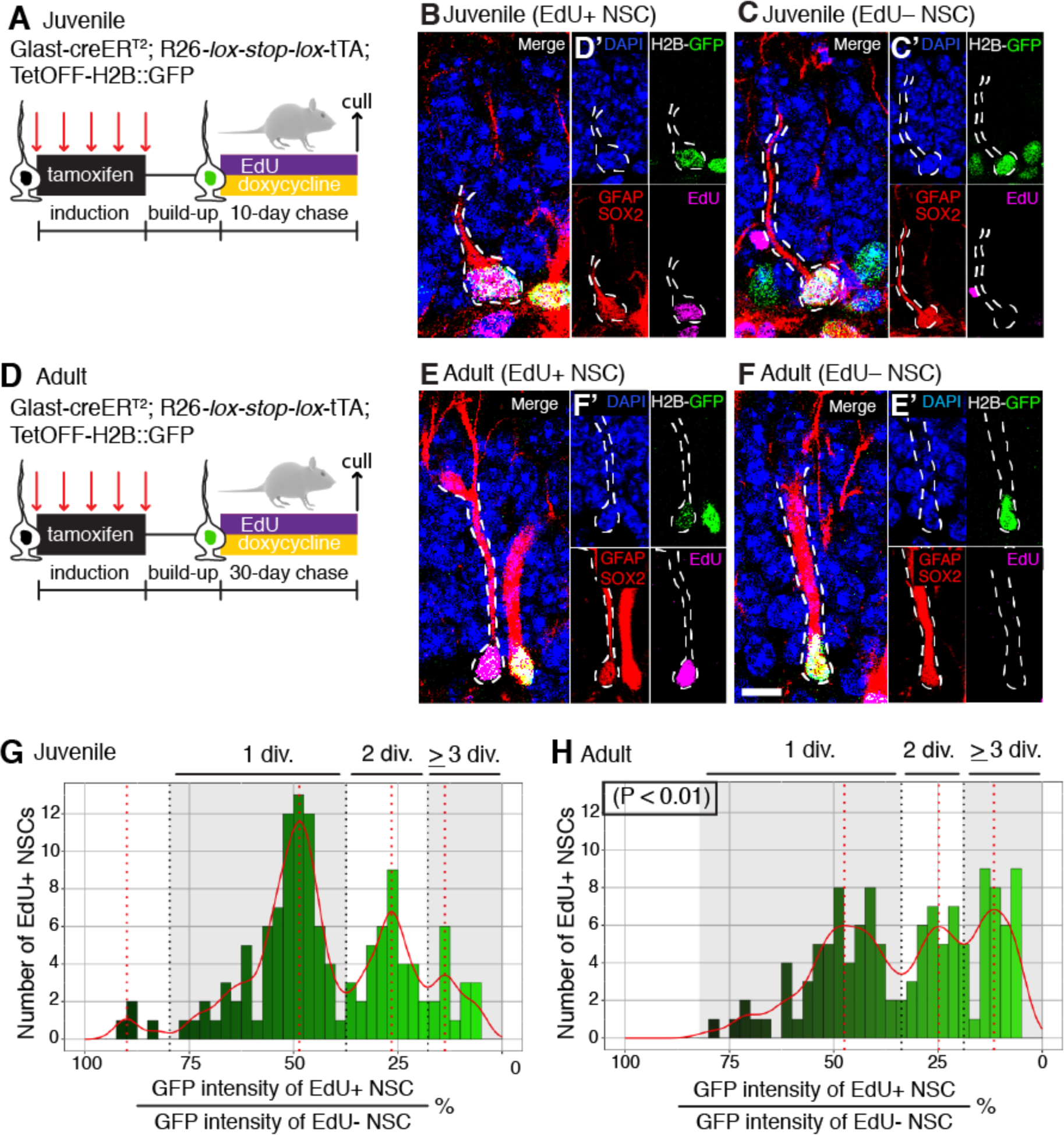
NSCs perform more self-renewing divisions with time. (**A**-**C**) A P14 cohort (juvenile) and (**D-F**) a 4.5-month-old cohort (adult) of *Glast-creER^T2^*; *R26-lox-stop-lox-tTA*; *TetOFF-H2B::GFP* mice were injected with tamoxifen to induce H2B::GFP expression. These juvenile and adult cohorts then received doxycycline for 10 or 30 days to stop label incorporation, respectively, and also received EdU to mark dividing cells. The chase lengths were adjusted to match the different length of time an activated NSC persisted before depleting in young and old mice (**B, E**) The H2B::GFP label became diluted in EdU+ NSCs but not (**C**, **F**) in quiescent EdU-NSCs. (**G**, **H**) The GFP signal from EdU+ NSCs was categorised into discrete bins through an automated analysis. These bins corresponded to the numbers of self-renewing divisions (0, 1, 2 and > 3 div.). EdU+ NSCs in (**H**) the adult mouse cohort showed increased label dilution than (**G**) in juvenile mice. Statistics: Fisher’s exact test to compare distributions in **G**, **H**. Abbreviations: Divisions (div.). Scale bar (in **F**): B, C, E, F = 15 μm, B’, C’, E’, F’ = 10 μm.

### Mathematical modelling of time-dependent changes

To support and extend these observations, we built a mathematical model of hippocampal NSC population dynamics at different ages. For model fitting, we used the numbers of total hippocampal NSCs, active NSCs and resting NSCs from mice 0.5-12 months of age (Figure 1A, B, I). In our model, dormant NSCs activate at the rate of R1. Once active, NSCs self-renew or deplete through terminal symmetric differentiation (we did not consider cell death as a major contributor to the loss of NSCs). Self-renewal is measured by the variable alpha (Marciniak-Czochra et al., 2009). After each self-renewing division (symmetric or asymmetric), the NSC passes through a resting state (G0) before re-activating at the rate of R2 (Figure S2A and Materials and Methods). In our dataset, we found that a decreasing activation rate provided a good fit for the numbers of total NSCs and active NSCs, as already found in a previous model of hippocampal NSC dynamics developed by Ziebell and colleagues (2018). However the model provided a poor fit to the quiescent label-retaining cell population (resting NSCs) since it did not take into account that dormant and resting cells could have different properties (Figure S2B). We therefore retained the decreasing rate of activation from the initial model but included more free parameters so that resting NSCs and dormant NSCs were allowed to have different activation rates, which resulted in a dramatic improvement of the fit of the model (AIC = 53 compared to 120 for the model where both populations have the same activation rate; Figure S2B). In the model of best fit (AIC = 36), resting NSCs across the first 6-months of life had a median activation rate 15-fold higher than that of dormant NSCs. Together, the accumulation of resting NSCs with a high activation rate and the decreasing activation rate of dormant NSCs with time help explain the overall increased contribution of resting NSCs to proliferation (Figure 2D). Altogether, the model we constructed supports the conclusion that the coordinated changes in behaviour of active, dormant and resting NSCs result in a decreased depletion of NSCs over time.

### Single cell RNA-sequencing and pseudotemporal ordering demonstrate state- and age-dependent changes to hippocampal NSC quiescence

The age-dependent changes in hippocampal NSC properties and in particular the different activation thresholds of resting and dormant NSCs warranted transcriptomic characterisation in order to identify the molecular mechanisms underlying these properties. To achieve this, we crossed the Nestin-GFP mouse line that labels NSCs (Mignone et al., 2004) with Ki67^TD^ mice to label proliferating cells and their progenies (Basak et al., 2018). The generated mice (hereafter, Ki67^TD-NES^) enabled lineage tracing of active NSCs. Ki67^TD-NES^ mice were injected with tamoxifen at 1-month, 2-months and 6-8 months of age to irreversibly label cells that were in *S-, G2-* and *M*-phase with red-fluorescence, their DG was then disassociated and GFP+ tdTomato- and GFP+ tdTomato+ cells were sorted by flow cytometry and sequenced together with the 10x Genomics platform (Figure 4A, B and Figure S3). We obtained a combined dataset of 24,189 cells (Table S1) which was integrated with Seurat (Stuart et al., 2019) and visualised with UMAP (Fig 4C) (Becht et al., 2018). The dataset was subsetted into NSCs and intermediate neuronal progenitors (IPCs) using marker genes *Hopx* and *Eomes* (Figure S4A, B) and then re-clustered to exclude IPCs and isolate only quiescent and active NSCs (Figure S4CD). From this final dataset of 2,947 cells, we used the trajectory inference tool Slingshot to infer progression along a pseudotime curve from quiescence to activation (Figure 4D, E) (Street et al., 2018). We validated this curve against an existing pseudotime analysis of 123 hippocampal progenitors containing NSCs and IPCs (Shin et al., 2015) by comparing genes statistically associated with progression in each dataset and found strong concordance between the two analyses (Figure 4F).

**Figure 4:**
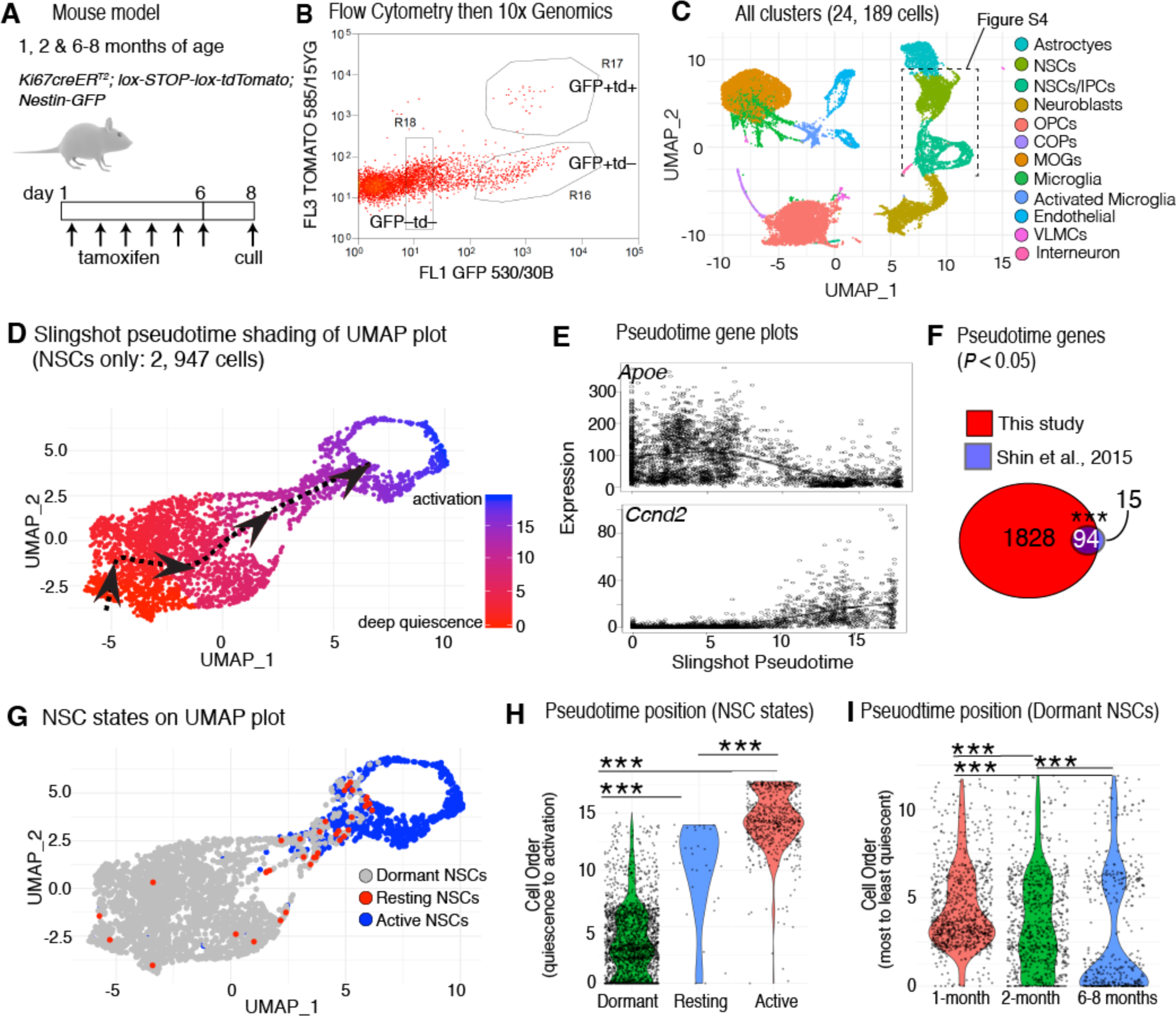
The quiescence depth of hippocampal NSCs is dependent on the proliferation history of the stem cell and the age of the mouse. (**A**) Cohorts of 1-, 2- and 6-8-month-old *Ki67-creER^T2^*; *lox-STOP-lox-tdTomato*; *Nestin-GFP* (Ki67^TD-NES^) mice were given tamoxifen and culled after 8 days, whereupon the DG was dissociated. (**B**) GFP+ tdTomato+ and GFP+ tdTomato-cells were collected by flow cytometry and sequenced using a 10x Genomics platform. (**C**) UMAP dimensional reduction showing the 24,189 cells sequenced from 8 experiments. (**D**) After two iterations of data sub-setting and re-clustering, a dataset of 2,947 NSCs was obtained and ordered using Slingshot, revealing a pseudotime trajectory from the most quiescent NSCs (red) to active NSCs (blue). (**E**) Pseudotime progression is negatively correlated with *Apoe* expression and positively correlated with *Ccnd2* expression. (**F**) Strong concordance between genes associated with this pseudotime trajectory and those from Shin. et al., 2015. Substantially more genes were statistically significant in our trajectory presumably due to increased dataset size. (**G**) UMAP plot showing the locations of dormant (quiescent and GFP+ tdTomato-), resting (quiescent and GFP+ tdTomato+) and active NSCs (proliferating and GFP+ tdTomato+/-). (**H**) Plotting pseudotime positions with violin plots reveals that resting NSCs are in a shallower state of quiescence than dormant NSCs and (**I**) that age increases the quiescence depth of dormant NSCs. Dots in plots represent individual cells. Statistics: Fisher’s exact test in **F**, Mann-Whitney U test in **H**, **I**. \*\*\**P*<0.001.

With this validated cell ordering, we analysed the positions along the pseudotime curve of NSCs in different states (active, resting and dormant) and of different ages. We identified active NSCs in G2/S/M-phase based on cell-cycle scoring algorithms (Butler et al., 2018). However, to distinguish active NSCs in G1 from quiescent NSCs in G0 among the recombined (tdTomato+) cells, we used ground truth testing (Figure S5) as no definitive criteria to distinguish G0 from G1 phase has yet been established (van Velthoven and Rando, 2019). We isolated bona-fide *S*-phase cells (based on cell-cycle scoring algorithms) and bona-fide G0 NSCs in a deep quiescence (based on their position in low-dimensional space; Figure S5A) and plotted the expression in these cells of genes known to be highly and stably expressed across both *G1-* and *S*-phases (i.e. *Mcm2-7*, *Ccne1/2*, and *Pcna*) (Braun and Breeden, 2007). This led to a clear separation of the S-phase and the G0-phase cell clusters, which allowed the definition of thresholds of gene expression (Figure S5B). These thresholds were then used to distinguish in the remainder of the dataset, tdTomato+ cells that were in G1 (that we marked as active NSCs) or in G0 (resting NSCs). Supporting our thresholding approach, resting NSCs expressed an independent set of G1/S-phase genes at similarly low levels as dormant cells (Figure S5C) and they had mRNA levels that were half those of active NSCs and equivalent to those of dormant NSCs, indicative of their quiescent state (Figure S5D).

As predicted from EdU labelling experiments (Figure 2), resting NSCs were rare (Figure 4G). They segregated along the pseudotime curve at positions intermediate between active NSCs and dormant NSCs (tdTomato- and quiescent). More than half of resting NSCs occupied the shallowest, least quiescent third of positions along pseudotime, whereas few resting cells occupied the deepest and most quiescent third of pseudotime positions (Figure 4G, H). Our single cell transcriptomics analysis therefore shows that the fraction of cycling NSCs that return to a state of quiescence are amongst the shallowest quiescent NSCs.

The effect of age on the depth of NSC quiescence was also striking (Figure 4I). The distribution of NSC pseudotime positions from 1-month-old mice was clearly overrepresented in lighter states of quiescence, while NSCs at 2-months of age were evenly distributed across the quiescence spectrum, and NSCs from 6-8-month-old mice were over-represented in deeper states of quiescence (Figure 4I). Thus hippocampal NSCs demonstrate an early and continuing progression into deeper quiescence. Altogether, the scRNA-seq data supports the EdU labelling and mathematical modelling data reporting the existence of a small population of resting NSCs that exist in a shallow quiescent state and the deepening quiescence of dormant NSCs.

### Division history affects the molecular properties of quiescent hippocampal NSCs

The previous findings from EdU-labelling experiments, mathematical modelling and pseudotime analysis of the single cell RNA-Seq data that resting cells have a higher activation rate and are in a shallower state of quiescence than dormant NSCs, led us to hypothesise that these two populations might diverge molecularly. To address this possibility, we used the scRNA-Seq data to compare gene expression in the different NSC populations. Gene ontological analysis of differential gene expression between resting and dormant NSCs (551 differentially regulated genes, Table S2), showed resting NSCs had reduced expression of genes associated with lipid metabolism (e.g *Apoe, Mt3, Prdx6* and *Acaa2*), suggesting a shallower quiescence as NSCs require *de novo* lipogenesis and fatty-acid oxidation to maintain quiescence (Knobloch et al., 2013; Knobloch et al., 2014) (Figure 5A). Resting NSCs also showed increased expression of genes associated with ribosomal biogenesis (e.g *Rpl17, Rpl5* and *Rpl29*), mRNA processing (e.g *Rbm3*, *Tra2b* and *Lsm7*) and translation elongation (e.g *Eef2* and *Eef1b2*), demonstrating an upregulation of protein biosynthetic pathways which has been associated with exit from quiescence (Figure 5B) (Cabezas-Wallscheid et al., 2017). Furthermore, multiple genes/pathways that have been associated with quiescence maintenance such as *Clu* (Basak et al., 2018), *Hopx* (Berg et al., 2019), *Notch2* (Engler et al., 2018) and *Id4* (*P* = 0.059) (Blomfield et al., 2019) were downregulated in resting NSCs compared with dormant NSCs (Figure 5A; Table S2). We validated the dataset by confirming that the expression of ID4 was downregulated at a protein level in resting versus dormant NSCs (Figure 5C-F). We extended this analysis to genes differentially expressed between resting and active NSCs. Excluding cell-cycle genes which were uniquely upregulated in active NSCs (Figure S5, Table S3), many of the aforementioned quiescence-specific genes were expressed in resting NSCs at an intermediate level between dormant and active states (Figure 5A, B and Table S3). Thus, although resting NSCs are quiescent, they show increased expression of metabolic and biosynthetic pathways and are thus already primed towards re-activation. This primed state (Llorens-Bobadilla et al., 2015) of resting NSCs demonstrates that the division history of a hippocampal NSC affects its dynamics and likely explains the increased contribution of resting NSCs to proliferation compared with dormant NSCs (Figure 2D).

**Figure 5:**
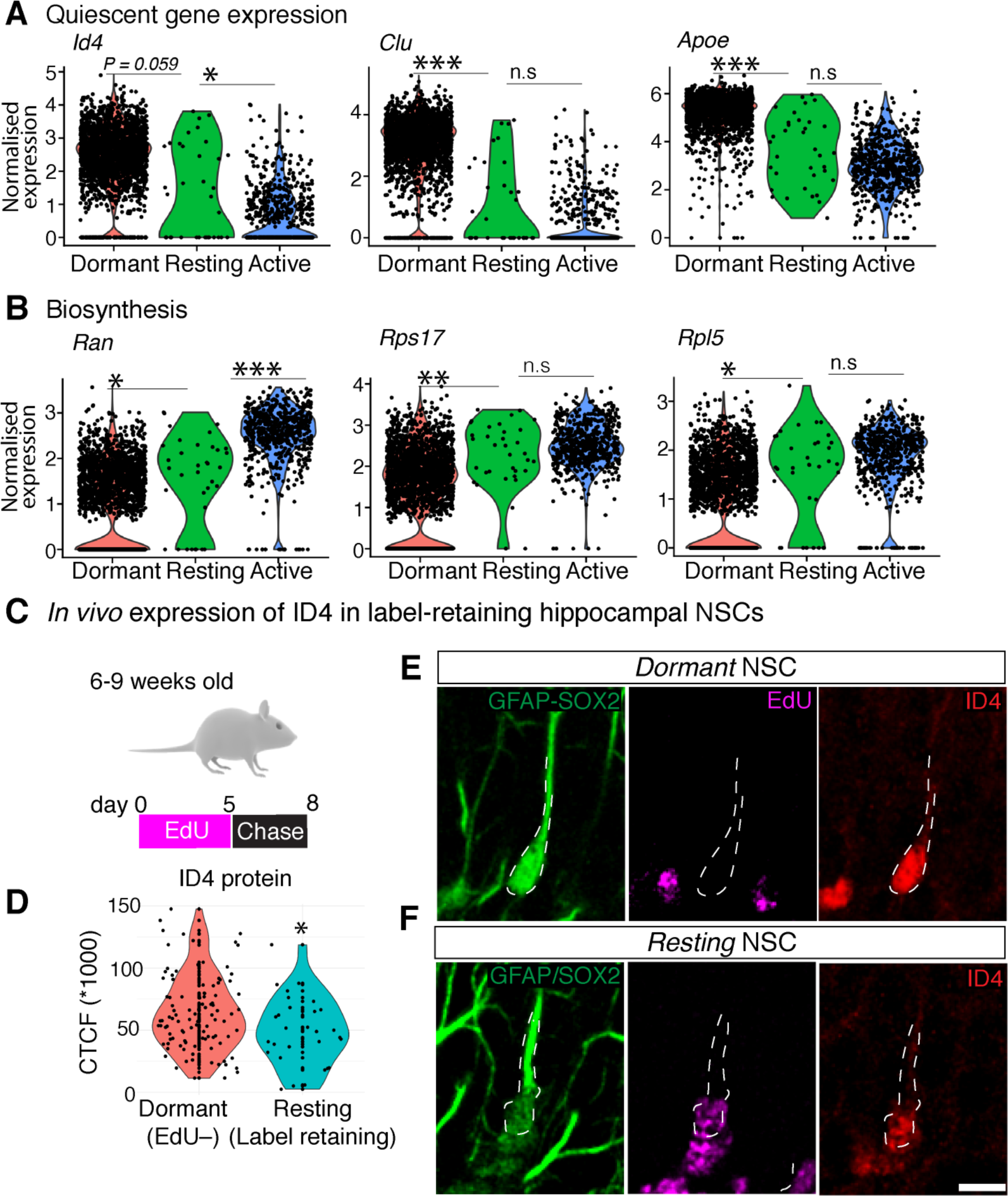
Resting NSCs display a distinct transcriptional profile indicative of a shallow/primed quiescent state. (**A**) Gene expression analysis from single cell transcriptomics data demonstrates that compared to dormant cells that have never activated, resting NSCs exhibit lower expression of quiescence marker genes and (**B**) increased expression of genes involved in biosynthesis, such as RNA transport and ribosomal biogenesis. (**C**) To validate the differential gene expression results, mice received EdU and were culled after a chase. (**D, F**) Label-retaining NSCs (EdU+, resting) had reduced immunolabeling of ID4 relative to (**D, E**) dormant NSCs (EdU-). Statistics:Student’s t-test performed on Pearson’s Residuals, with FDR corrected *P*-value in **A**, **B**. Student’s t-test in **D**. *\*P<0.05, ** P <* 0.01, *\*\*\*P<0.001*. Scale bar (in **F**): **E, F** = 10 μm.

### Declining ASCL1 levels explain time-dependent changes in quiescence

The fact that NSC properties change coordinately over time, with dormant NSCs entering a deeper state of quiescence as active NSCs increasingly return to a resting state and undergo greater numbers of self-renewing divisions, suggested that a common molecular mechanism might underlie these different changes. The transcription factor ASCL1 was a strong candidate to contribute to the changing properties of NSCs, as induction of ASCL1 expression is required for the activation of quiescent NSCs (Andersen et al., 2014), a reduction of ASCL1 protein levels by degradation by the E3 ubiquitin ligase HUWE1 allows cells to return to a resting state (Urban et al., 2016), and a reduction in ASCL1 levels increases the likelihood of embryonic NSCs self-renewing rather than differentiating (Imayoshi et al., 2013). We reasoned that the changing properties of hippocampal NSCs might reflect progressively lower expression levels of ASCL1 over time.

*Ascl1* is transcribed by most NSCs (Blomfield et al., 2018). However, because of low protein levels due to degradation induced by HUWE1 (Urban et al., 2016) and by ID factors (Blomfield et al., 2019), ASCL1 protein cannot be detected with anti-ASCL1 antibodies in quiescent NSCs and can only be seen when ASCL1 protein levels are at their highest, i.e. in active NSCs. To increase ASCL1 detection sensitivity, we examined its expression in a mouse line that expresses an ASCL1-VENUS fusion protein (Imayoshi et al., 2013). The labelling of the fusion protein with anti-GFP antibodies greatly enhanced the detection of ASCL1 compared with an anti-ASCL1 antibody, so that low levels of ASCL1 were readily detectible even in quiescent hippocampal NSCs (Figure S6). Consistent with our hypothesis, there was a progressive but dramatic reduction in the proportion of NSCs (including quiescent NSCs) that express detectable levels of ASCL1-VENUS throughout adulthood (Figure 6A, B). 54% of NSCs expressed ASCL1-VENUS in 0.5-month-old mice, 19% in 2-month-old mice, and 3% in 12-month-old mice (Figure 6C). There was a parallel reduction in the mean ASCL1-VENUS fluorescence intensity of NSCs in 6-month old mice relative to 1-month old mice (Figure 6D). Thus, the observed changes in hippocampal NSCs properties with age indeed correlate with a reduction in ASCL1 protein levels in NSCs.

**Figure 6:**
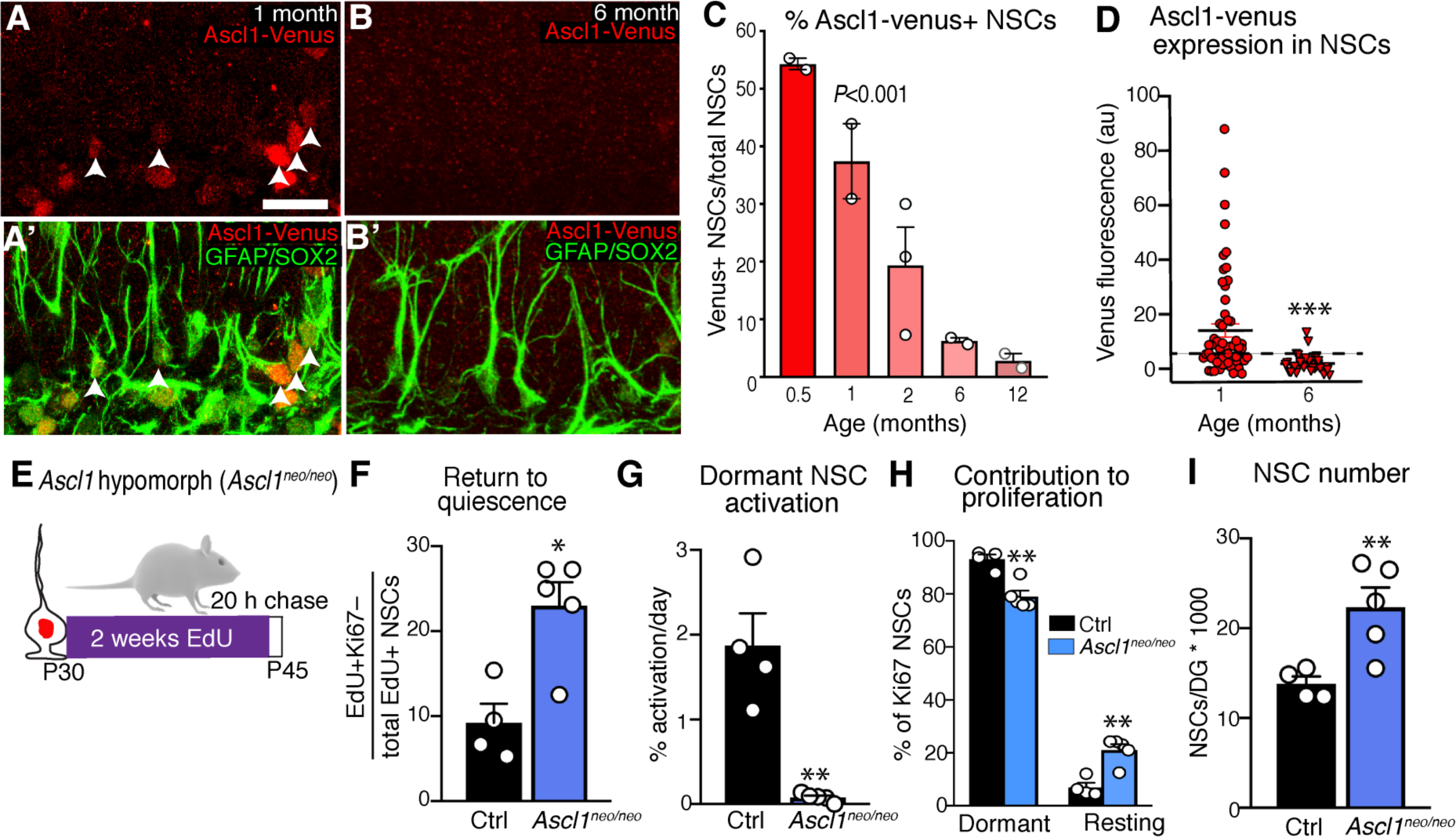
Ascl1 protein levels decrease with time and cause progressive changes to NSC behaviour. (**A**, **B**) Immunolabeling of NSCs (GFAP+ SOX2+) that express the ASCL1-VENUS fusion protein in the DG of (**A**, **A’**) 1-month- and (**B**, **B**’) 6-month-old mice using a sensitive anti-GFP antibody. Arrowheads indicate ASCL1-VENUS+ NSCs. (**C**) Fewer NSCs are positive for ASCL1-VENUS with age (**D**) and the mean expression intensity of ASCL1-VENUS fusion protein decreases. (**E**) *Ascl1*^neo/neo^ and control mice were given EdU to label active and resting NSCs and were culled following a chase. (**F**) Hippocampal NSCs in *Ascl1*^neo/neo^ mice were increasingly returning to quiescence after activating (EdU+ Ki67-) and (**G**) were activating much less from a dormant NSC state than in control mice. (**H**) Resting NSCs contributed more to the proliferative NSC pool in *Ascl1*^neo/neo^ mice than in controls and (**I**) had more hippocampal NSCs. Graphs represent the mean ± s.e.m. Dots represent individual mice, except **D** where dots represent individual cells (minimum 20 cells analysed per mouse, 2 mice per age). Statistics: One-way ANOVA in **C**, Student’s t-test in **D**, **F-I**. \**P*<0.05, ** *P* < 0.01, *\*\*\*P<0.001*. Scale bar (in **A**): **A**-**B** = 21.9 μm.

We had previously shown that deleting *Ascl1* from NSCs in 2-month-old mice stops the depletion of hippocampal NSCs, which demonstrated a link between ASCL1 expression and NSC depletion (Andersen et al., 2014). To establish causality, we utilised an Ascl1 hypomorphic mouse line (*Ascl1*^neo/neo^) that expresses reduced levels of *Ascl1* in adult hippocampal NSCs (Andersen et al., 2014). We labelled proliferating, resting and dormant NSCs in 1-month-old *Ascl1*^neo/neo^ mice by EdU retention and Ki67 labelling as before (Figure 6E). Compared to age-matched controls, EdU+ NSCs in hypomorphic mice were 2.5 times more likely to return to quiescence (Figure 6F) and dormant NSCs were 24 times less likely to exit quiescence (Figure 6G). As a result, resting NSCs contributed more to the proliferative pool in *Ascl1*^neo/neo^ mice than in controls (Figure 6H) and the total number of NSCs was increased, presumably due to a lower depletion rate (Figure 6I). Therefore, hippocampal NSCs in *Ascl1*^neo/neo^ mice precociously resemble NSCs from older mice, which provides direct evidence that the declining levels of ASCL1 drive the time-dependent changes in hippocampal NSCs behaviours and the resulting reduction in NSC depletion rate.

### Declining ASCL1 levels are due to increased post-translational degradation

In our scRNA-seq analysis of hippocampal NSCs in 1-month-old and 6-month-old mice, the levels of *Ascl1* transcripts did not change significantly over time (Figure 7A). We therefore reasoned that the steep time-dependent decline in ASCL1 protein levels must be due to altered post-translational regulation, such as through increased expression or activity of ID proteins or the E3 ubiquitin-ligase HUWE1, which both promote the targeting of ASCL1 for proteasomal destruction (Blomfield et al., 2019; Urban et al., 2016). Indeed, the expression of *Huwe1* increased in 6-month-old NSCs relative to its expression in 1-month-old NSCs (Fig 7B), while among *Id* genes, *Id4* was upregulated but *Id1-3* were downregulated with age (Table S4). We therefore focused on the role of *Huwe1* in driving the time-dependent change in ASCL1 protein expression. The HUWE1-dependent degradation of ASCL1 has been shown to promote the return of hippocampal NSCs to a resting state (Urban et al., 2016) and it also suppresses the activation of dormant NSCs (Figure S7A, B). Thus, the increased expression of *Huwe1* and/or increased efficiency of HUWE1-mediated ASCL1 degradation might contribute to the progressive reduction in ASCL1 levels, the subsequent changes in NSC properties and the long-term maintenance of the NSC pool.

**Figure 7:**
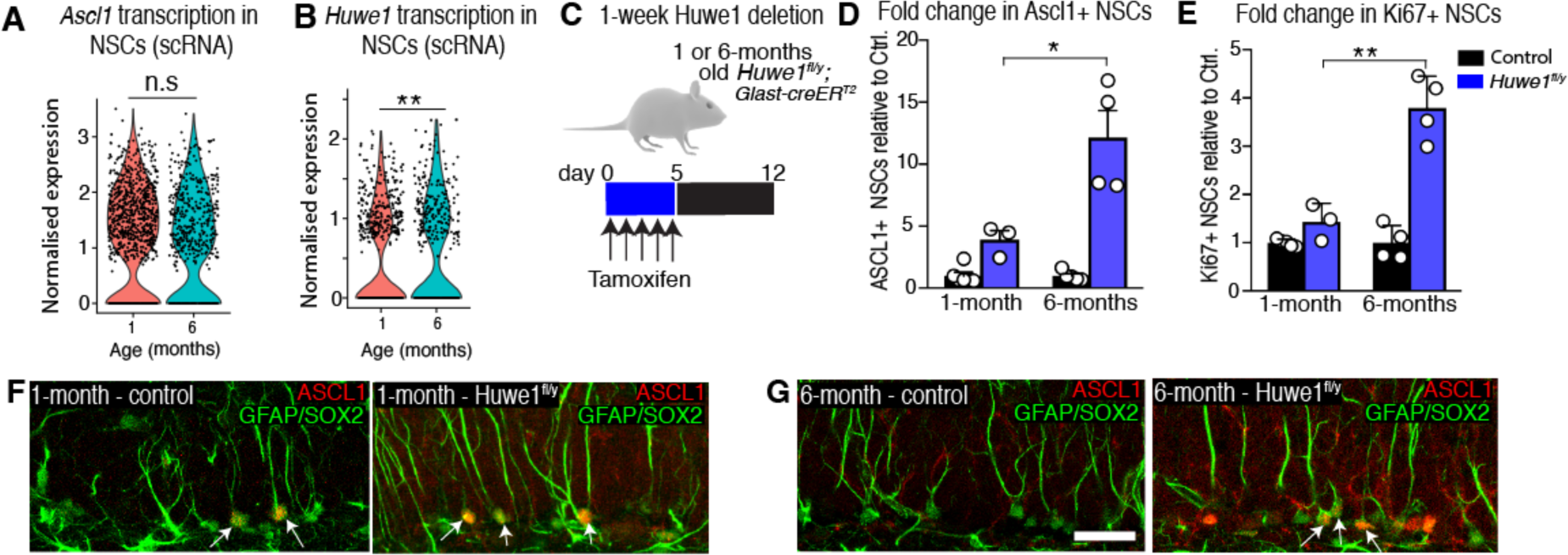
Expression and activity of Huwe1 increases with time. (**A**) Gene expression analysis from single cell transcriptomics data shows that *Ascl1* transcript levels in NSCs are largely unchanged between young and adult mice, (**B**) while expression of the ubiquitin-ligase *Huwe1* increases. (**C**) 1- and 6-month-old *Huwe^fl/y^* and control mice were culled 1-week after receiving tamoxifen. (**D, E**) The numbers of ASCL1+ and Ki67+ NSCs were increased upon *Huwe1* deletion in both 1- and 6-month-old mice but the effect was much larger in the older cohort. Images of control and *Huwe^fl/y^* mice at (**F**) 1-month and (**G**) 6-months of age, 1-week after tamoxifen, with arrows indicating Ascl1+ NSCs. Graphs represent mean ± s.e.m. Dots represent the mean of individual mice. Statistics: Student’s t-test performed on Pearson’s Residuals, with FDR corrected *P*-value in **A**, **B**, Student’s t-test in **D** and **E**. \**P*<0.05, ** *P <* 0.01. Scale bar (in **G**): **F, G** = 28 μm.

To test whether HUWE1 activity increases with age, we deleted *Huwe1* from hippocampal NSCs by administering tamoxifen to *Huwe1*^fl/y^; *Glast*-*creER^T2^* mice at 1- and 6- months of age and analysed the mice 1 week later (Figure 7C). As expected, the deletion of *Huwe1* elicited an increase in the number of NSCs expressing ASCL1. However this increase was substantially larger in 6-month-old mice (12.1-fold increase) than in 1-month-old mice (3.9-fold increase) when normalised to age-matched controls (Figure 7D and Figure S7D).

Similarly, the number of Ki67+ NSCs increased by a larger amount in 6-month-old mice (3.8 fold) than in 1-month-old mice (1.4 fold) (Figure 7E and Figure S7E), indicating that Huwe1 activity, which eliminates ASCL1 expression and suppresses proliferation of hippocampal NSCs, is greater in 6-month-old than in 1-month-old mice. Altogether, these results demonstrate that the expression and activity of *Huwe1* increase with time to reduce ASCL1 protein levels and mediate the associated changes in NSC quiescence that serve to preserve the NSC pool.

## DISCUSSION

It is currently unclear whether human hippocampal neurogenesis extends beyond childhood (Moreno-Jimenez et al., 2019; Sorrells et al., 2018). Learning about the mechanisms that control the rate of neurogenesis in other species may shed light on changes that occur or perhaps fail to occur in humans, to preserve lifelong neurogenesis. In most mammals, high numbers of new neurons are generated in the hippocampus in juveniles, their numbers decline rapidly in young adults and are then maintained at low levels throughout adulthood and late in life (Amrein, 2015; Ben Abdallah et al., 2010; Snyder, 2019). In mice, the rapid decline in neurogenesis in juveniles and young adults reflects a rapid reduction in numbers of NSCs, which has been ascribed to self-consuming divisions whereby NSCs divide to produce only neurons or astrocytes and no new NSCs (Encinas and Sierra, 2012; Gontier et al., 2018; Pilz et al., 2018). In this study we demonstrate that the relative maintenance of NSCs throughout adulthood is due to the decoupling of NSC activation from depletion, leading to the emergence of a subpopulation of resting NSCs that sustain proliferation. These resting NSCs exist in a shallow quiescence and their emergence results in increased self-renewal capacity. In contrast, dormant NSCs that have not previously divided become more deeply quiescent, providing a reserve of NSCs that infrequently activate under physiological conditions. This series of changes are caused by a progressive reduction in the levels of the pro-activation protein ASCL1 due to increased rates of post-translational degradation.

The changes in NSC behaviours we identify serve to preserve the NSC pool and long-term neurogenesis and are therefore distinct from stem cell exhaustion that occurs during aging. Indeed, most of the changes we describe occur in the first 6 months of life when mice are of breeding age, suggesting an adaptive function. Aging is defined by functional decline and the accumulation of cellular damages resulting in loss of fitness (Lopez-Otin et al., 2013). In contrast, we found that the reduction of NSC number during this period is not associated with a loss of NSC function. Instead, the numbers of self-renewing divisions increases, explaining previous reports that the proportion of hippocampal NSCs that are actively dividing at any given time remains largely stable (Encinas et al., 2011; Ziebell et al., 2018). This is in contrast to the V-SVZ niche where the active NSC fraction decreases substantially. Interestingly, the increased quiescence of NSCs in the V-SVZ niche was found to occur as early as 7-months of age and to be mediated through changes in inflammatory signalling. It was suggested that this early change might not reflect stem cell dysfunction but help preserve the NSC pool (Kalamakis et al., 2019). Our transcriptional analysis of NSCs in 1-month-old and 6-month-old mice shows little change over time in inflammatory, mitochondrial and lysosomal pathways that typify stem cell aging (Table S4) (Leeman et al., 2018). Instead, we propose that hippocampal NSCs progress during adulthood through a process of extended development. The late developmental processes experienced by NSCs may be comparable to changes that occur between “fetal” and “adult” hematopoietic stem cells, which differ in their self-renewal capacities, turnover rates and differentiation potentials (Copley et al., 2013), with the difference that changes in hematopoietic stem cells occur more abruptly, between 3 and 4 weeks of age in mice, 1 and 2 years in baboons and 1 and 3 years in humans, while the changes that we report in NSCs are more protracted and span the juvenile and much of the adult phases.

A central finding of this study is that proliferating hippocampal NSCs increasingly return to a distinct, shallower state of quiescence with time. Multiple lines of evidence support the conclusion that the resting state of NSCs truly represents quiescence and not merely slow divisions, including 1) the emergence of non-dividing, EdU label-retaining NSCs that remain in the hippocampal niche for extended periods that can exceed 30 days; 2) a subset of Ki67-CreER-traced NSCs analysed by scRNA-seq that lack expression of cell cycle genes and cluster between the bulk quiescent NSC population and active NSCs when using dimension reduction techniques; 3) these resting NSCs contain levels of mRNA similar to those of dormant NSCs and half those of active NSCs, which is a characteristic of quiescence. Additional evidence supports the resting state as a distinct, shallower state of quiescence than that of dormant NSCs, including, 4) that resting NSCs express quiescence-associated transcription factors and metabolic genes at levels intermediate between those of dormant and active NSCs; 5) that resting NSCs have a greater rate of activation than dormant cells, and 6) that they contribute disproportionately to the proliferating NSC pool although their population only represents a small fraction of all quiescent NSCs even at advanced ages.

Adult tissue stem cell compartments often have several stem cell populations that differ in their level of quiescence (Barker et al., 2010). Previous experiments have demonstrated that NSCs in the adult niches of the V-SVZ and DG lie upon a trajectory from quiescence to activation, including a population of primed NSCs in the V-SVZ (Llorens-Bobadilla et al., 2015; Shin et al., 2015). However, whether each NSC operates along the same trajectory, whether the trajectory can be reversed, and whether subpopulations of NSCs along this trajectory can be linked to specific cellular behaviours, i.e. returning to quiescence, remained unclear. Our scRNA-seq dataset, which contains approximately 10-20 fold more adult hippocampal NSCs than the largest previous study (Hochgerner et al., 2018), and includes tracing with the *Ki67-creER^T2^* allele allowing for the unbiased sampling of active NSCs unlike lines featured in other studies (Ibrayeva et al., 2019; Pilz et al., 2018), reveals two distinct quiescent states. We find in particular that hippocampal NSCs that have recently divided and returned to a resting state are in much shallower quiescent state than NSCs that have never left quiescence. Although resting NSCs present lower expression levels of quiescence-associated transcription factors and higher levels of metabolic genes than dormant NSCs, they nevertheless are able to remain quiescent due to an active degradation of the pro-activation factor ASCL1 that prevents NSC activation and cell cycle re-entry (Urban et al., 2016). However, the reduced expression of quiescence-associated genes maintains resting NSCs in shallow quiescence or primed state which allows them to reactivate much more readily than dormant cells and thereby sustain NSC proliferation and neurogenesis over the long term. Resting NSCs likely belong to a larger population of primed cells, including cells that are transiting from dormancy to activation (Llorens-Bobadilla et al., 2015). However our characterisation of resting NSCs uniquely demonstrates how this primed state can arise based on the proliferative history of each hippocampal NSC. Moreover, whether activated dormant NSCs transiting through a primed state and proliferating NSCs returning to a resting state reach functionally and molecularly identical or divergent primed states, remains to be addressed.

Proliferating NSCs that return to a transient resting state rather than differentiating have additional opportunities to divide, which likely explains the increased number of self-renewing divisions that NSCs undergo over time. Whether resting NSCs re-activate and return to quiescence once or multiple times is unclear and long term live-imaging studies (Pilz et al., 2018) should resolve this question. The finding of increased self-renewal does not contradict the results of our mathematical model. In our ranking of different mathematical models each with their own permutations and free parameters, a model similar to the best fit model but predicting increased self-renewal was placed second overall only because it contained this additional free parameter (AIC = 41 vs 36; Figure S2E), suggesting that increased self-renewal might have less impact on age-related changes compared to other mechanisms. In a previous modelling study, Ziebell and colleagues (2018) found that with age, hippocampal NSCs spent more time in quiescence between divisions and concluded that NSCs were more likely to (re)activate in old mice. However this model did not consider NSC heterogeneity and specifically that NSCs that have returned to a resting state have a higher activation rate than NSCs that have always remained quiescent. Our new model accounts for how NSC behaviour is altered after activation and return to a resting state, and supports the idea that the increased prevalence of resting NSCs contributes to the slowing down of the rate of NSC depletion, the maintenance of NSC proliferation and the extension of hippocampal neurogenesis into old age.

The deepening quiescence over time of dormant NSCs, which have not activated since the establishment of the DG niche, fits well with findings from *in vitro* studies of fibroblasts indicating that the depth of cellular quiescence is proportional to the time spent in quiescence (Kwon et al., 2017; Soprano, 1994). The transcriptional changes associated with a deeper quiescent state are typically an exaggeration of the changes that occur initially upon quiescence induction (Coller et al., 2006). In this study, many of the genes that were downregulated in dormant *versus* active hippocampal NSCs at 1-month of age, indeed showed further reductions at 6-months of age, including for example, in the expression of ribosomal genes. This progression into deeper quiescence did not result in mis-regulation of pathways associated with pathological features of aging, emphasising the difference between a deep quiescent state and senescence (Kwon et al., 2017). The deepening quiescence of dormant NSCs may instead enable these cells to act as a reserve pool of stem cells, suggesting that these cells should not necessarily be considered as targets to enhance tissue resiliency (Ibrayeva et al., 2019). Eventually, some fraction of dormant NSCs could become so deeply quiescent so as to become irreversibly arrested, i.e. senescent, and contribute to the pathological features of aging (Martin-Suarez et al., 2019) but this seems unlikely to occur in mice that have only recently become sexually mature.

Our data suggests that current conflicting models of hippocampal NSC behaviour can be reconciled by considering changes in behaviour over time. We find in juvenile mice that NSCs uniformly adhere to the disposable stem cell model (Encinas et al., 2011). Thereafter, heterogeneity emerges so that some hippocampal NSCs acquire the capacity to return to quiescence and self-renewal increases, as reported in Bonaguidi et al. (2011). Importantly, the age of the mice used in the original studies might have contributed to their divergent conclusions. Two-month-old mice were used in the BrdU-labelling experiments that originated the disposable stem cell model (Encinas et al., 2011), and 10-week-old mice were used when live-imaging led to similar conclusions (Pilz et al., 2018). Three to four-month-old mice were used in the labelling experiments where a subpopulation of hippocampal NSCs were found to return to quiescence (Urban et al., 2016), while a substantial part of the clonal experiments that posited long-term self-renewal was analysed in 1-year-old mice (Bonaguidi et al., 2011). Altogether, these observations suggest that in addition to technical differences (Bonaguidi et al., 2012), the use of mice of different ages contributed to the development of contradictory models of hippocampal NSC behaviour. The effect of time we describe here renders these models compatible.

We have recently reported that the pro-activation gene *Ascl1* is broadly transcribed by most quiescent NSCs, though the protein is detected only in active NSCs using conventional antibody labelling techniques, due to destabilisation by ID4 and subsequent proteasomal destruction in quiescent NSCs (Blomfield et al., 2019). Here, we demonstrate that in young mice and when using sensitive antibodies against a VENUS fusion protein, ASCL1 protein is found broadly and at low levels in many quiescent NSCs and that these levels are gradually reduced over months during juvenile and adult stages. These findings mirror a recent report showing that pancreatic epithelial progenitors express high levels of the *Neurog3 m*RNA but low and often undetectable levels of the NGN3 protein, which nonetheless played an important role in maintaining the progenitor state (Bechard et al., 2016). We credit the changes in hippocampal NSCs dynamics throughout adulthood to the declining levels of ASCL1 protein. Reduced ASCL1 levels should decrease the activation rate of dormant NSCs since ASCL1 is required for NSC activation (Andersen et al., 2014). Moreover, the reduction in ASCL1 levels by accelerated protein degradation has been shown to increase the probability of cells leaving the cell cycle and entering the resting state (Urban et al., 2016). We examined juvenile *Ascl1* hypomorphic mice and observed indeed a deeper quiescence of dormant NSCs and an accelerated accumulation of resting NSCs, indicating that the precocious reduction of ASCL1 expression in these animals produces cellular changes normally only seen later, during adulthood. NSCs also undergo greater numbers of self-renewing divisions in adult than juvenile mice. In embryonic NSCs, low levels of ASCL1 have been associated with reduced differentiation and increased self-renewal (Imayoshi et al., 2013), which might contribute to the additional rounds of NSC divisions and resulting label dilution observed in adult mice compared with juveniles.

We found that the reduction in ASCL1 protein levels was likely due to increased post-translational degradation since *Ascl1* mRNA was largely unaffected by age. Indeed both the expression and activity of the ubiquitin-ligase *Huwe1*, which targets ASCL1 for proteasomal destruction (Urban et al., 2016), increased throughout adulthood. The mechanisms leading to the transcriptional upregulation of *Huwe1* and/or its increased activity are unclear. It has been shown that the phosphorylation status of substrates can affect HUWE1-mediated ubiquitination rates (Forget et al., 2014). Interestingly, ASCL1 stability has been shown to be phospho-dependent (Ali et al., 2014) and we identified a number of phosphatases and kinases that are differentially expressed in NSCs from adults versus juveniles (Table S4). Identifying niche signals that control the phosphorylation status of ASCL1 and its interaction with HUWE1 might provide insights into timing mechanisms that drive changes in hippocampal NSC behaviour during adulthood. Altogether, our results demonstrate how a series of early, progressive and coordinated changes in hippocampal NSC properties function to preserve proliferation and thereby neurogenesis beyond the juvenile period in mice. Whether similar mechanisms are present in humans to extend neurogenesis beyond childhood warrants investigation.

## AUTHOR CONTRIBUTIONS

Conceptualization [LH, TS, AMC, FG], Data curation [LH], Formal analysis [LH, PR, ZG, TS], Validation [LH, PR, TS], Investigation [LH, PR, SA, MDMM, TS], Visualization [LH, PR, TS], Methodology [TS, NU], Writing—original draft [LH, FG], Project administration [FG], Writing-review and editing [LH, PR, TS, ZG, SA, MDMM, AE, NU, AMC, FG]. Supervision [FG], Funding acquisition [FG], Project administration [FG].

## ACKNOWLEDGMENTS

We gratefully acknowledge Lan Chen for technical support, Rekha Subramaniam, Nicholas Chisholm and Michael Nagliati for managing the mouse colony; the Advanced Sequencing Facility, Bioinformatics and Biostatistics Facility, Experimental Histopathology Facility and Flow Cytometry Facility of the Francis Crick Institute for their help and support; Ana Martin-Vivalba for providing feedback on manuscript; Ryoichiro Kageyama for providing mice. This work was supported by the Francis Crick Institute, which receives its funding from Cancer Research UK (FC0010089), the UK Medical Research Council (FC0010089) and the Wellcome Trust (FC0010089). This work was also supported by the UK Medical Research Council (project grant U117570528 to FG), by the Wellcome Trust (Investigator Award 106187/Z/14/Z to FG) and the German Research Foundation DFG (SFB 873; subproject B08, Anna Marciniak-Czochra).

## METHODS

### LEAD CONTACT AND MATERIALS AVAILABILITY

Further information and requests for reagents should be directed to and will be fulfilled by the Lead Contact, Francois Guillemot.

### EXPERIMENTAL MODEL AND SUBJECT DETAILS

Wild-type animals in this study were on a C57Bl/6J genetic background (000664, The Jackson Laboratory). In contrast, because most transgenic mice had multiple transgenes, these were maintained on a mixed genetic background, with littermates serving as controls. Mice with conditional *Huwe1* (Zhao et al., 2008), *Ascl1*^neo/neo^ (Andersen et al., 2014), *R26tTa* (Wang et al., 2008), *tdTomato* (Madisen et al., 2010) and *YFP* (Srinivas et al., 2001) alleles have been described before; as have the cre-driver lines *Glast*-creER^T2^ (Mori et al., 2006), *Ki67*-*creER^T2^* (Basak et al., 2018) and the reporter lines *Ascl1*-*venus* (Imayoshi et al., 2013), *Nestin-GFP* (Mignone et al., 2004) and *H2B*::*GFP* (Kanda et al., 1998). Both males and female mice were used throughout the study; an exception to this was for the x-linked *Huwe1* conditional allele, where only males were used.

All experimental protocols involving mice were performed in accordance with guidelines of the Francis Crick Institute, national guidelines and laws. This study was approved by the UK Home Office (PPL PB04755CC). Throughout the study, mice were housed in standard cages with a 12 h light/dark cycle and *ad libitum* access to food and water.

### METHOD DETAILS

#### Mathematical modelling

Details of mathematical modelling are found in data file 1.

#### Preparation and sectioning of mouse brain tissue

Mice were perfused transcardially with phosphate buffered saline (PBS) or saline (5-10 mL), followed by 4% paraformaldehyde (10-20 mL) and post-fixed for 16-24 h before long term storage in PBS with 0.02% sodium azide, at 4°C. Brains were sectioned in a coronal plane at 40 μm using a vibratome (Leica, Deerfield, IL). The entire rostral-caudal extent of the hippocampus was collected in a 1 in 12 series.

#### Antibodies, immunofluorescence and cell counts

At least 1 series per mouse (5-6 sections), per experiment, was stained and analysed. For experiments requiring the detection of fixation-sensitive antigens (ASCL1, TBR2, Ki67, MCM2) free-floating sections were subjected to heat-mediated antigen retrieval in sodium citrate solution (10 mM, pH 6.0) at 95°C for 10 min. A shorter duration of heat-retrieval was used (2-5 min) if the detection of GFP was required, which is heat sensitive (Nakamura et al., 2008). Following incubation at 95°C, sections were cooled for 5 min before being rinsed with PBS and blocked in 2% normal donkey serum diluted in PBS-Triton X-100 (0.1%) for 2 hours. Sections were then incubated overnight at 4°C with primary antibodies diluted in blocking buffer, washed 3 times in PBS for 10 min, and incubated with secondary antibodies at room temperature for 2 h. Sections were stained with 4’,6-diamidino-2-phenylindole (DAPI, 1:10K, Thermo Fisher Scientific) and mounted onto Superfrost slides (Thermo Fisher Scientific) with Aqua PolyMount (Polysciences).

The stained sections were imaged using a 40X oil objective on a SP5 or SP8 Leica confocal microscope through a depth of 15-20 μm, with a step size of 1 μm (X/Y pixel diameter = 0.28-0.38 μm). Cell counts were then performed on the imaged sections. To present normalised data in units of cells/DG, the area of the sampled SGZ was measured (SGZ length*z-stack depth). The calculated surface area was then scaled to an age-appropriate reference dataset from C57Bl/6J mice of different ages (0.5, 1, 2, 6, 12 and 18 months) where the total SGZ surface area had been calculated.

#### Tamoxifen treatment and administration of EdU

Tamoxifen solution (10-20 mg/mL) was prepared for intraperitoneal injections by dissolving the powder (T5648, Sigma, St Louis, MO) in a mix of 10% ethanol and 90% cornflower oil (C8267, Sigma). The dose and injection regime were selected according to the relative difficulty of inducing recombination of the floxed allele, whereas the route of administration changed from intraperitoneal injection to oral gavage dosing for experiments that lasted longer than 5-days, based on veterinary advice.

For experiments involving the *Huwe1^fl^* lines, mice were injected with 80mg/kg of tamoxifen per day, whereas the daily dose delivered to *Ki67*^TD^ mice in Figure S1 and *H2B-GFP* mice in Figure 3, was 100mg/kg. Finally, for the Ki67^TD-NES^ mice used in the scRNA-seq experiments they were delivered tamoxifen via oral gavage for 6 consecutive days at 100mg/kg. In all experiments, control mice received tamoxifen at the same doses of experimental mice. The control mice were littermates that were wildtype for the floxed alleles and contained the *cre* transgene or were homozygous for the floxed alleles but negative for *cre*.

The labelling of cells progressing though S-phase was performed by either intraperitoneal injections of EdU (50 mg/kg) or through administration of EdU in the drinking water (0.2 mg/ml). For long-term EdU water administration, fresh solution was replaced at least every 56 hours. The sensitivity of EdU detection kits allowed for lower concentration of EdU to be used than in standard BrdU drinking water assays (typically 1mg/mL), and avoided obvious cellular toxicity (Young et al., 2013).

#### Calculation of observed *versus* expected NSC depletion rate (disposable stem cell model)

The observed NSC depletion rate was measured by first quantifying the reduction in NSC number between two timepoints (for example, between 0.5 months and 1 month of age). This was converted to the rate of change per day, which was normalised to the size of the NSC pool at the earlier timepoint (i.e. 0.5 months). This was repeated for all timepoints (2, 6, 12 and 18 months). The expected rate of depletion predicted by the disposable stem cell model was calculated by reducing the observed depletion date at the earliest instance (between 0.5-1 months of age) by the percentage reduction in the size of the active NSC pool for all later timepoints (2, 6, 12 and 18 months).

#### Calculation of contribution to NSC proliferation

The contribution of dormant and resting NSCs to the proliferative (Ki67+) NSC pool is reported in Figure 2 and 6 from experiments in which mice received EdU in the drinking water for 2-4 weeks, prior to 20 h chase period and culling (Figure 1F, Figure 2A). The number of Ki67+ NSCs originating from the resting NSC pool was determined by multiplying the fraction of Ki67+ cells that were EdU+, with the probability that the EdU+ cell had returned to a quiescence at some stage during the EdU exposure (see Figure 1E for probability of return to quiescence). The fraction of Ki67+ NSCs originating from the dormant NSC pool was determined by subtracting the contribution of resting NSCs from the total proliferative pool.

#### H2B-GFP label dilution experiments

The dilution of the H2B-GFP nuclear label was used as a proxy to determine the number of self-renewing NSC divisions. Juvenile (0.5 months) and adult (4.5 months) cohorts were injected with tamoxifen (100mg/kg), each day for 5 days, to activate the CreER^T2^ recombinase, recombine the *R26tTa* locus and induce H2B-GFP expression. In the adult cohort, the H2B:GFP label was allowed to build up for 25-days after tamoxifen administration. To ensure the juvenile cohort remained relatively young by the end of the experiment, the H2B:GFP was allowed to build up for just 14-days after tamoxifen administration, which was sufficient to produce bright and even labelling of NSC nuclei. After H2B-GFP labelling, the juvenile and adult cohorts were given 2 mg/ml doxycycline in drinking water, with EdU (0.2mg/mL) and sucrose (1% w/v) for 10 and 30 days, respectively, switching off H2B-GFP expression. The solution was changed 3-times per week. The length of this label-dilution period was adjusted between ages to match the approximate lifespan of an EdU+ NSC in young and old mice. The juvenile cohort received a shorter treatment, because EdU+ NSCs rapidly divide and differentiate, such that >90% have differentiated after 10 days (Figure 1K). Conversely, in 6-month-old mice, it takes 30-days for >90% of the EdU+ NSC population to differentiate (Figure 1K). After doxycycline treatment, juvenile mice (now 6-weeks of age) and adult mice (now 6-7 months of age) were immediately perfused and processed for immunostaining.

Sections were stained using antibodies against GFAP and SOX2 and were also stained for EdU and DAPI. Cell counts and GFP fluorescence intensity measurements were performed on the entire DG at different levels along the rostrocaudal axis. The GFP fluorescence intensity of EdU+ NSCs was compared to the mean GFP fluorescence intensity of 2-3 neighbouring EdU-NSCs, to calculate a ratio. To minimise variability caused by staining or imaging artefacts, the EdU-cells that were selected for comparison had to be in the same X-Y image tile (291*291 μm) and their centre of mass within 4 z-planes (3.52 μm) of the EdU+ NSC. Segmentation of nuclei was performed using the Fiji plugin ‘3D Object Counter’ (Bolte and Cordelieres, 2006), and the raw integrated density for each z-level was summed and used for the measurement output. Data points were excluded if the standard error of the mean of the average fluorescence intensity of EdU-NSCs was greater than 15% of the mean, as inclusion of this data detrimentally affected normalisation.

The R package “multimode” (Ameijeiras-Alonso et al., 2018) was used to estimate the most probable number of clusters (modes) in each dataset (with the “modetest” function) and to locate the modes and anti-modes (by means of the “locmodes” function). The modes and anti-modes were then used to create bins that separated the values based on the number of cell divisions ! that single EdU+ NSCs performed and the number of values falling in each bin was counted for each dataset. Therefore, we obtained the fraction of cells that divided !-times in each dataset.

#### Single-cell analysis of Ki67^TD^; Nestin*-*GFP mice

##### DG dissection, FAC sorting and 10x Chromium

Each scRNA-seq experiment comprised 2-5 male and female Ki67^TD^; Nestin*-*GFP mice that had been injected with tamoxifen for 6 consecutive days (100mg*kg) and culled on the 8^th^ day. In total, 1 experiment was performed with 1-month-old mice, 3 experiments with 2-month old mice, and 3 experiments with 6-8-month-old mice (Table S1).

The protocol for microdissection of the DG was performed as described (Hagihara et al., 2009; Walker and Kempermann, 2014). The dissected DG was then disassociated using the Neural Tissue dissociation kit (P) (Milteny Biotec, 130-092-628) according to manufacturer’s instructions, with the following exceptions: a 37°C orbital shaker was used during the enzymatic digestions, and we used manual trituration with fire-polished pipettes to aid dissociation following the incubations with enzymatic mix 1 and 2. The dissociated cells were centrifuged and resuspended in 750μl recovery media (0.5% PBS-BSA in DMEM/F12 without phenol red (Sigma, DG434) and 1ug/ml DAPI). The cells were sorted on the MoFlo XDP (Bechman Coulter) using a 100μm nozzle with a pressure of 30 psi, and a sort efficiency of more than 80%. The events were first gated to remove debris (fsc-h vs ssc-h), to remove aggregates (typically pulse-width vs area then pulse-height vs area/width) and to remove dead cells (fsc-h vs DAPI fluorescence). Cells were then gated for tdTomato expression according to a control mouse that expressed Nestin-GFP alone; whereas the GFP gate was set according to the expected distribution of GFP fluorescence intensities based on prior control experiments. Cells were collected in 700 μl of recovery media in 1.5mL tubes, and spun down at 500G for 7 min at 4°C. After removing all but 50μl of the supernatant, the cells were then gently resuspended using a wide-bore pipette. The single-cell suspension (to a maximum of 10,000 cells) was then loaded into the 10x Chromium.

##### Sequencing and mapping

We prepared one library for each of the 8 samples across the 7 experimental days. The libraries for 2-month-old mice were prepared with 10x Genomics Chemistry, Single Cell 3’ version 2, while the 1-month and 6-month libraries were prepared with version 3 (Table S1). After sequencing, cellRanger count (Ver. 3.0.2) was used to map the FASTQ files to our custom mouse genome (mm10-3.0.0), which contained 4 elements. The first element was the GFP coding sequence to detect transcription of Nestin-GFP. The second, was the 5’ floxed sequence of the *tdtomato* allele that detects transcription of the intact *tdtomato* locus. Finally, we added 3’ regions of the *tdtomato* locus that are transcribed upon recombination, and encode the *woodchuck hepatitis virus post-transcriptional regulatory element* (WPRE) and the *bovine growth hormone* polyA signal (bGH-polyA). The custom genome was made with the cellranger mkref command.

##### Seurat analysis: Quality control

The cellranger aggr command was used to merge the count files of the 9 individual libraries without normalisation. The merged matrix was then read into Seurat (Ver. 3.1.2) for analysis (Stuart et al., 2019). GEM beads were kept for further analysis if they contained more than 500 genes and fewer than 10% mitochondrial reads in order to remove GEM beads that contained only background signal or a dead cell. Doublets/multiplets were then removed based on the expression of more than 1 marker gene per GEM bead (e.g. co-expression of the astrocytic marker *Aldoc* with the oligodendrocyte marker *Mog)* In total, 24,169 of 25,990 (94.5%) passed these quality control steps. Later, during the first subsetting/re-clustering of the data (visualisation of the NSCs and IPCs alone) three further small doublet clusters appeared, which were identified and removed based on the lack of specific marker expression (Briggs et al., 2018).

We then added metadata that marked each cell for tdTomato expression and cell-cycle status. Cells were identified as tdTomato+ if they contained < 4 reads of the intact *tdTomato* locus, and more than > 1 read of the recombined *tdTomato* locus based on ground-truth testing of our first dataset, in which we sequenced the tdTomato+ and tdTomato-populations separately (Figure S3). We also identified whether a cell was in Go or actively cycling based using ground-truth testing. We isolated bona-fide S-phase cells through the CellCycleScoring function in Seurat, and bona-fide Go-phase NSCs based on UMAP position. We examined genes highly expressed across G1 and S-phase *(Mcm2-7, Ccne1/2,* and *Pcna),* binarized each cell as positive or negative for each marker and plotted every cell according to their index score (range of possible values 0-9). These plots revealed a clear separation of two cellular populations which we used to define thresholds for G0/G1. Specifically, cells in which we detected > 1 cell-cycle gene were scored as in G1, while all other cells were considered quiescent/post-mitotic. Cells sequenced with Version 3 of the 10x Chemistry had on average twice the number of UMIs detected, so the thresholds for tdTomato and cell-cycle scoring were doubled.

##### Seurat analysis: Data visualisation

The merged Seurat object was split according to library/experimental day and the data transformed using the SCTransform function. The 5,000 most variable genes and anchors across the 8 datasets were then identified and the data integrated using the default parameters. The integrated dataset was then visualised using UMAP (Becht et al., 2018). Elbow plots were used to determine the number of significant principal components. Following cluster identification with known marker genes, we extracted all NSCs and IPCs and re-clustered these cells using the same workflow. We iterated this process a further time to then extract only NSCs, this time splitting and integrating the Seurat object according to 10x Chemistry (V2 or V3), as this best eliminated technical variation without removing biological signal (assessed by clustering of replicates across libraries). We appended metadata classifying these cells as dormant NSCs (quiescent and tdTomato-), resting NSCs (quiescent and tdTomato+) or active NSCs (active and tdTomato+/-). The final dataset contained 2,947 cells which we processed for differential gene expression and pseudotime ordering.

##### Pseudotime ordering

To construct a pseudotime ordering of NSCs we used the trajectory inference tool Slingshot (Street et al., 2018) (Ver. 1.2.0) and UMAP dimensionality reduction. The curve generated by Slingshot was used to plot the pseudotime position of each cell between conditions and to generate a set of genes statistically associated *(P* < 0.05) with pseudotime progression. For the reconstruction of the pseudotime trajectory from Shin and colleagues (2015) we read in the normalised counts (TPM) from GSE71485 into Seurat. Next PCA analysis was performed using the 5 top components and the data was then visualised using UMAP. We excluded a cluster of cells that lacked significant expression of *Hopx* that largely corresponded to contaminating oligodendrocytes, as reported in the original study. We compared the set of genes that were statistically associated with pseudotime with our dataset.

### QUANTIFICATION AND STATISTICAL ANALYSIS

#### Single-cell data

Differential gene expression analysis was performed using the FindMarkerGene function in Seurat using the Pearson residuals located in the “scale.data” slot of the SCT assay using Student’s t-test (Hafemeister and Satija, 2019). For each comparison, genes were only tested for differential expression if they were expressed by a minimum of 20% of cells in at least one of the two conditions being compared. No minimum log fold change threshold was enforced. Genes were considered as statistically significant if they had FDR adjusted *P*-value of < 0.05.

#### Statistical analysis of cell counts

The statistical testing approach were specified prior to data collection and implemented using Graphpad Prism (version 8.0) or in R (3.6.2). Two-tailed unpaired Student’s t-tests were performed when comparing two groups. For experiments involving one or two independent variables, two-way ANOVA was performed and the *P*-values from two-way ANOVA are reported in the text. Any significant main effect of genotype or age detected by two-way ANOVA was followed by multiple t-tests using a pooled estimate of variance where appropriate, and significance was corrected for using the Holm-Sidak method in Prism 8.0 (Graphpad). Details of statistical tests and sample size for specific experiments are found in figure legends.

## DATA AND CODE AVAILABILITY

Upon publication data and code will be deposited in suitable public databases.

## SUPPLEMENTAL ITEMS

**Data file 1, related to** Figure 1**-3 and Figure S2:** Detailed description of mathematical model.

**Table S2, related to** Figure 5: Differential gene expression analysis of resting *versus dormant* hippocampal NSCs

**Table S3**, r**e**lated **to** Figure 5: Differential gene expression analysis of resting *versus active* hippocampal NSCs

**Table S4, related to** Figure 6**, 7:** Differential gene expression analysis of quiescent hippocampal NSCs from 6-month *versus* 1-month mice

## SUPPLEMENTAL ITEMS

**Figure S1, related to Figure 1:**
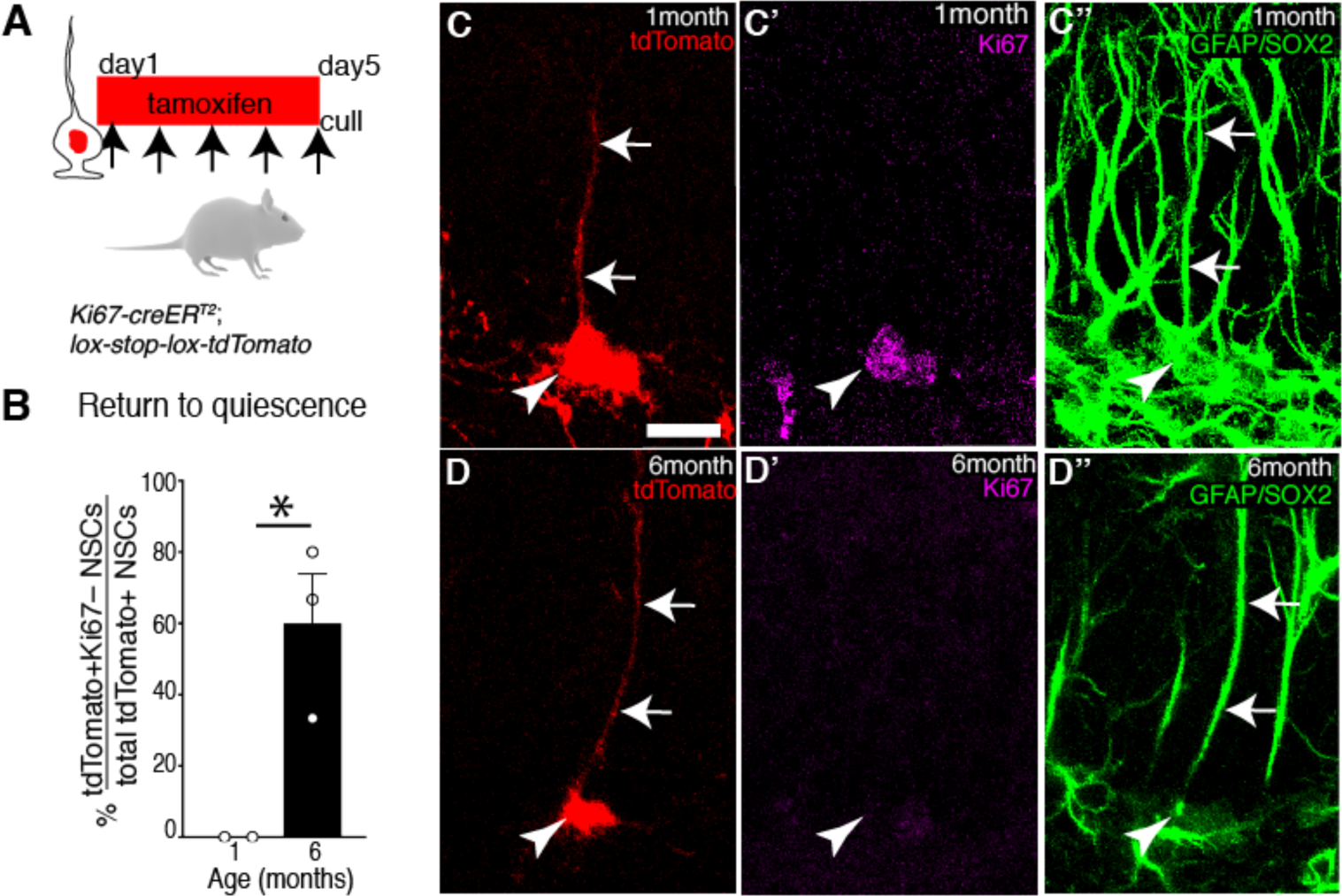
Genetic lineage tracing of active NSCs reveals they increasingly return to quiescence with time. (**A**) Upon tamoxifen administration, active NSCs and their progeny, including resting NSCs, are indelibly labelled with tdTomato fluorescence in *Ki67-creER^T2^*; *lox-STOP-lox-tdTomato* (Ki67^TD^) mice. (**B**, **C**) In 1-month-old mice culled immediately after tamoxifen treatment, tdTomato+ NSCs remain active (Ki67+), whereas in 6-month-old animals (**B**, **D**) a substantial proportion returns to quiescence (Ki67-). In (**C**, **D**) arrowheads indicate NSC cell body, arrows indicate the radial tdTomato+ process. Graph in **B** shows mean ± s.e.m, dots represent individual mice. Statistics: Student’s t-test in **B**. Scale bar (in **C**): **C**, **D** = 15 μm. \**P*<0.05.

**Figure S2, related to Figure 1-3:**
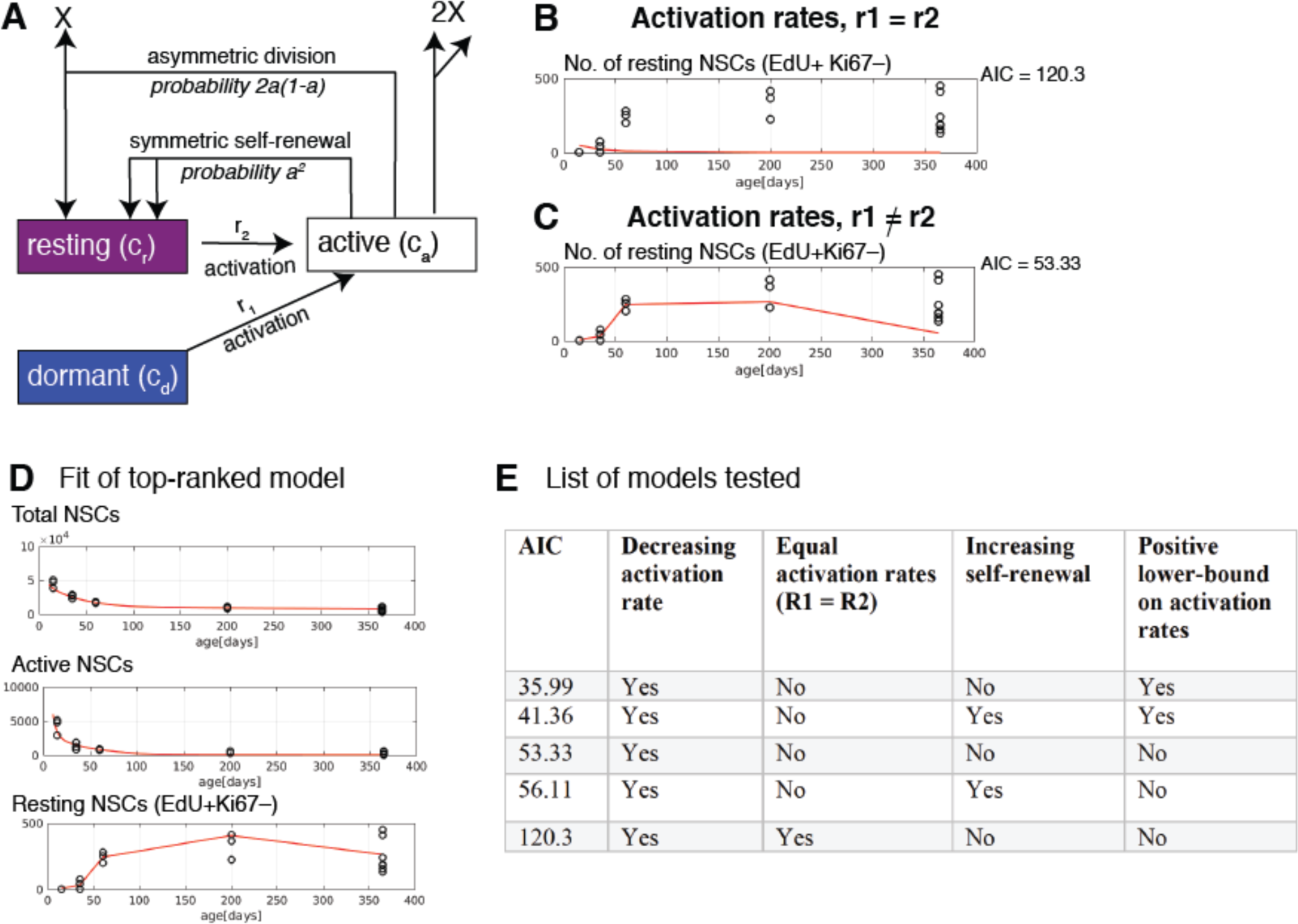
Mathematical modelling of time-dependent changes. (**A**) Development of a mathematical model favoured the scenario where (**B**, **C**) dormant and resting NSC activation rates decreased with time but were higher for resting than for dormant NSCs, as shown in the fit of these models to the number of resting NSCs against time. (**D**) Fit of top model (lowest AIC) to number of dormant NSCs, active NSCs and resting NSCs. (**E**) List of models investigated. (AIC = Akaike Information Criterion).

**Figure S3, related to Figure 4:**
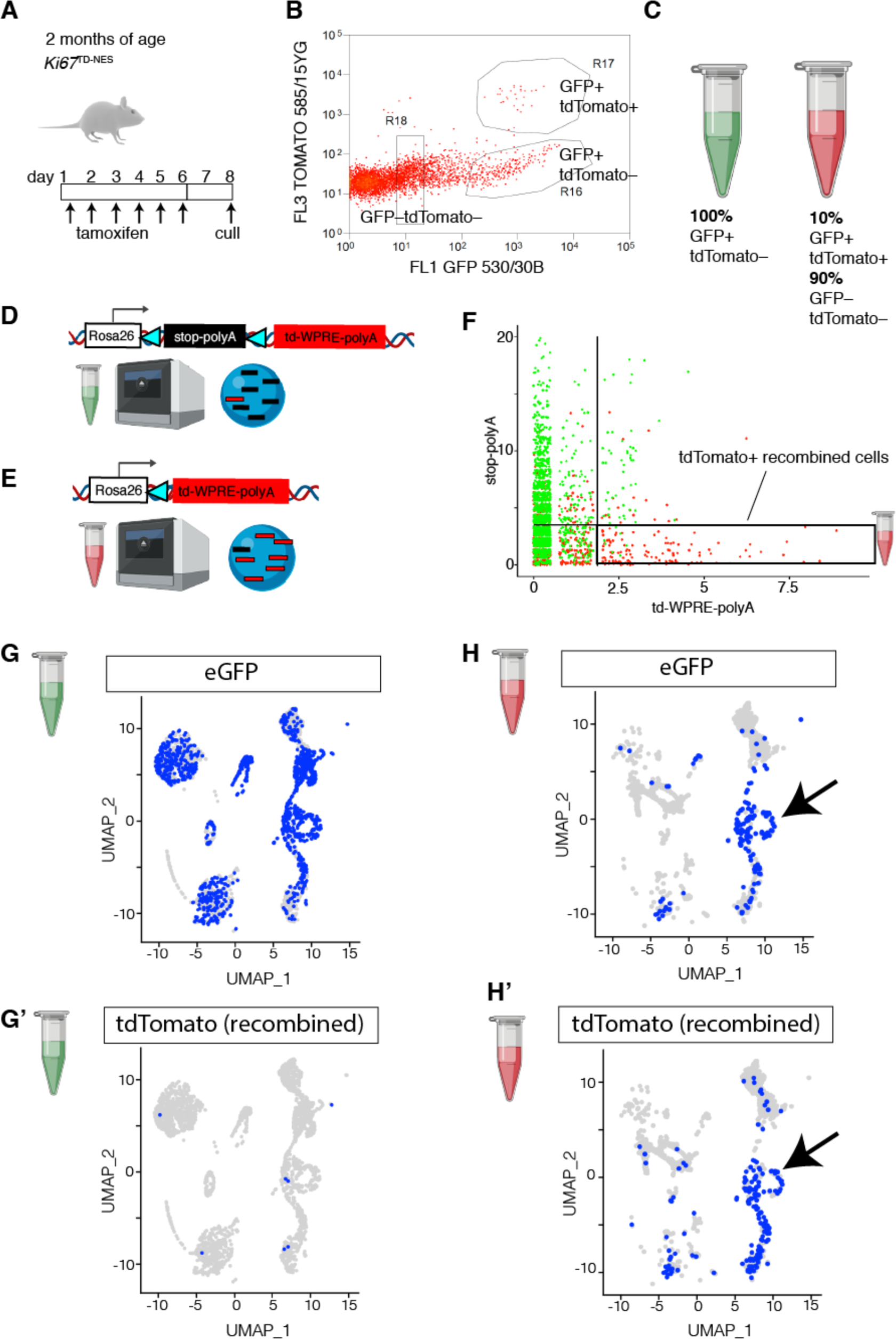
Post-sequencing identification of cells that have undergone *cre*-mediated recombination in Ki67^TD-NES^ mice. (**A**) Schematic of pilot experiment to determine the efficacy of identifying tdTomato+ cells after sequencing in *Ki67-creER^T2^*; *lox-STOP-lox-tdTomato*; *Nestin-GFP* mice (Ki67^TD-NES^). (**B**, **C**) GFP+ tdTomato-cells and GFP+ tdTomato+ cells were collected in separate tubes. Because GFP+ tdTomato+ cells were rare (∼1,000 in this experiment) they were also mixed with 9,000 GFP-tdTomato-cells to ensure there was a minimum of 10,000 cells to centrifuge and recover before loading into 10x Genomics. (**D**, **E**) The two tubes were loaded in 10x machine. (**D**) Recombined cells should have low/absent expression of the floxed “*td*-*stop*::polyA” transcript, whereas (**E**) they should have higher expression of the downstream “*td::WPRE::bGH*::polyA” transcript. (**F**) Scatterplot of intact *versus* recombined transcripts demonstrates that cells with a recombined signature almost entirely (95.67%) come from the GFP+ tdTomato+ tube. (**G**) UMAP plots of cells from GFP+ tdTomato-tube demonstrate almost all cells are GFP+ and tdTomato-(only 2.68% are tdTomato+) (**H**) UMAP plots of cells from GFP+ tdTomato+/GFP-tdTomato-tube shows that in contrast, approximately 90% of the cells are negative for both markers, corresponding to the GFP-tdTomato-cells that were loaded to weight the GFP+ tdTomato+ sample, whereas the remaining cells have high expression of GFP and tdTomato and localise to proliferative cells, i.e. intermediate neuronal progenitors and neuroblast clusters (arrow).

**Figure S4, related to Figure 4:**
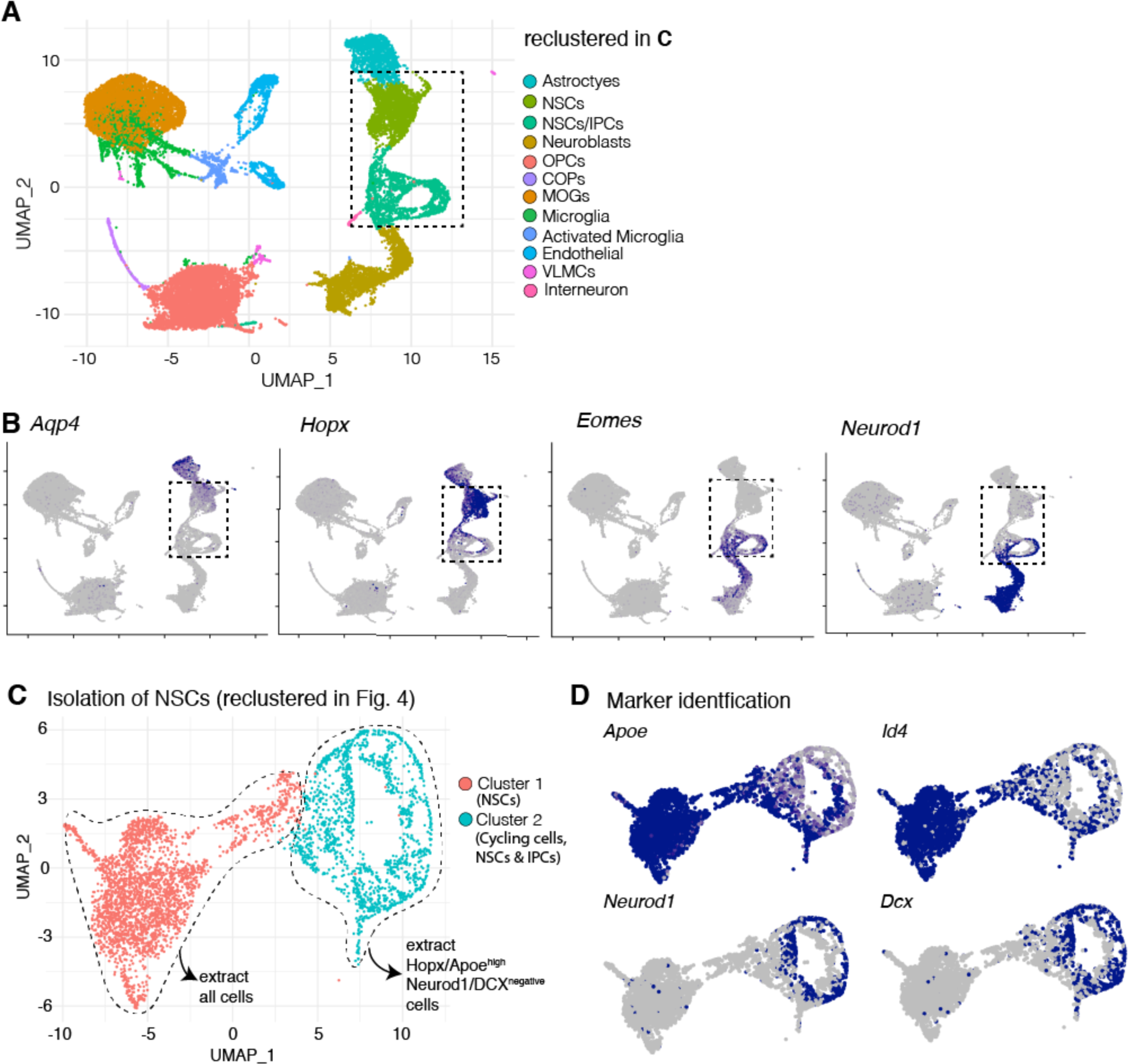
Identification and isolation of NSCs from merged dataset. (**A**) UMAP plot of cell clusters from merged dataset of 8 libraries (different ages and replicates) with their cellular identity indicated with the colour key. (**B)** The NSC cluster was identified based on an *Aqp4*-low and *Hopx*-high expression profile. The IPC cluster was identified based on an *Eomes*-high and *Neurod1*-high expression profile. Dashed boxes in **A** and **B** indicate the selection and isolation of NSC and IPC clusters for re-clustering in **C**. (**C**) UMAP plot displaying NSCs and IPCs from merged dataset. Cluster 1 was comprised entirely of NSCs and cluster 2 was comprised of cycling cells (IPCs and some NSCs). All cells in cluster 1 were extracted as NSCs for further analysis, (**D**) whereas only cells that had high *Hopx*/*Apoe* expression and were negative for both IPC *mRNA* markers *Neurod1* and *Dcx* were extracted as NSCs from cluster 2. These extracted cells were then re-clustered and are presented in Figure 4D, G.

**Figure S5, related to Figure 5:**
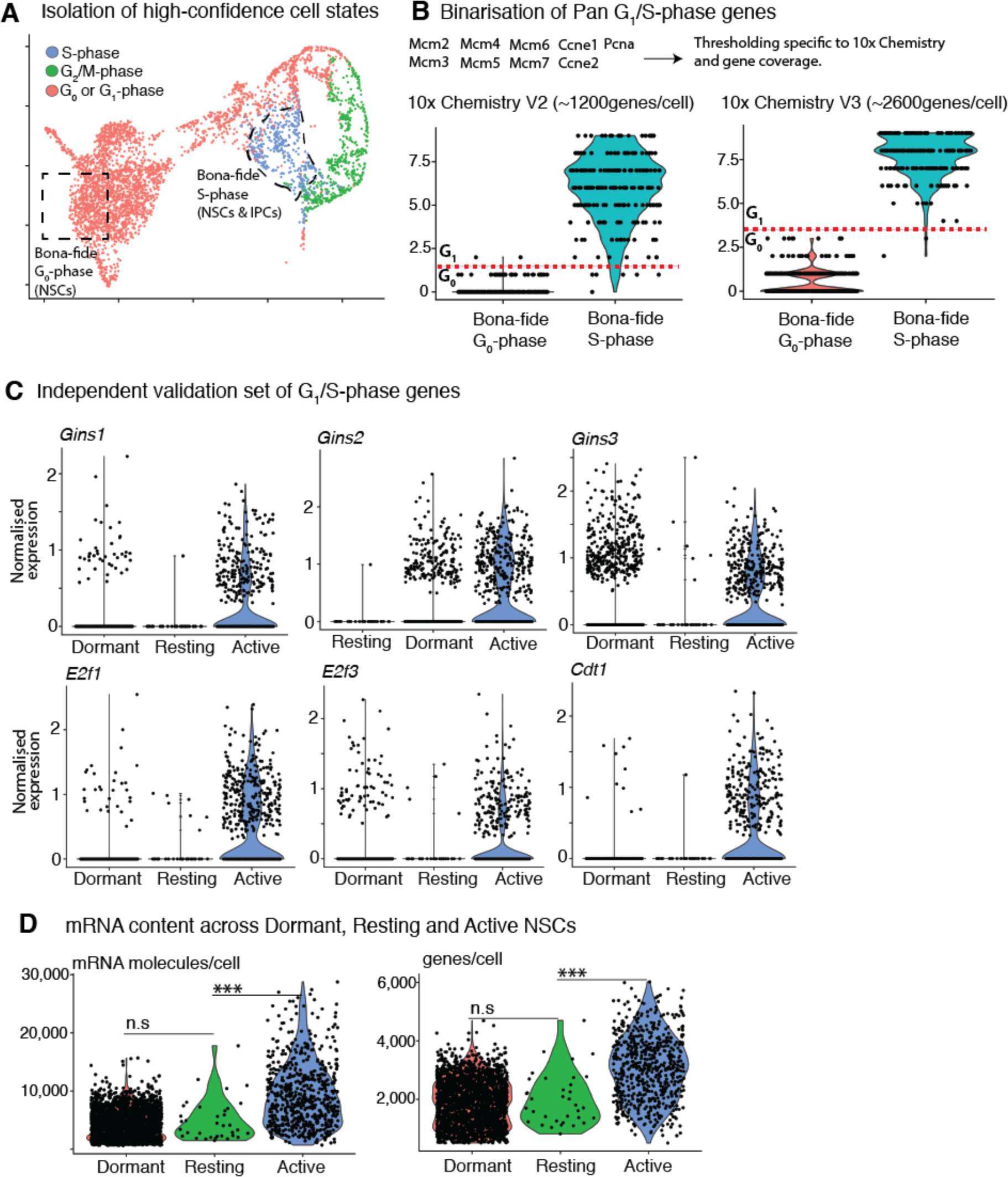
Thresholding criteria to distinguish G0 from G1 phase. (**A**) UMAP plot showing NSC and cycling cell (NSC/IPC) clusters. For ground-truth testing bona-fide S-phase NSC/IPCs were identified through the CellCycleScoring function in Seurat, whereas bona-fide G0-phase NSCs were isolated based on position in UMAP plot corresponding to deep quiescence. (**B**) Genes that are highly and stably expressed in both G1- and S-phase were binarized and added together, to generate an index. Violin plots of this index showed a clear hourglass separation between G0 and S-phase cells, which allowed for thresholds (separate thresholds for each version of the 10x kit) to be chosen to distinguish G0 and G1 in the remainder of the dataset. Cells above the threshold were designated as G1 and therefore as actively cycling cells alongside S- and G2/M-phase cells. (**C**) This thresholding was tested in an independent set of G1- and S-phase genes, which showed that resting NSCs do not express significantly higher levels of these genes than dormant NSCs, and much lower levels than active NSCs. (**D**) Attesting to the quiescent state of resting NSCs they also have half the level of mRNA as active NSCs, comparable to dormant NSCs. \*\*\**P*<0.001.

**Figure S6, related to Figure 6:**
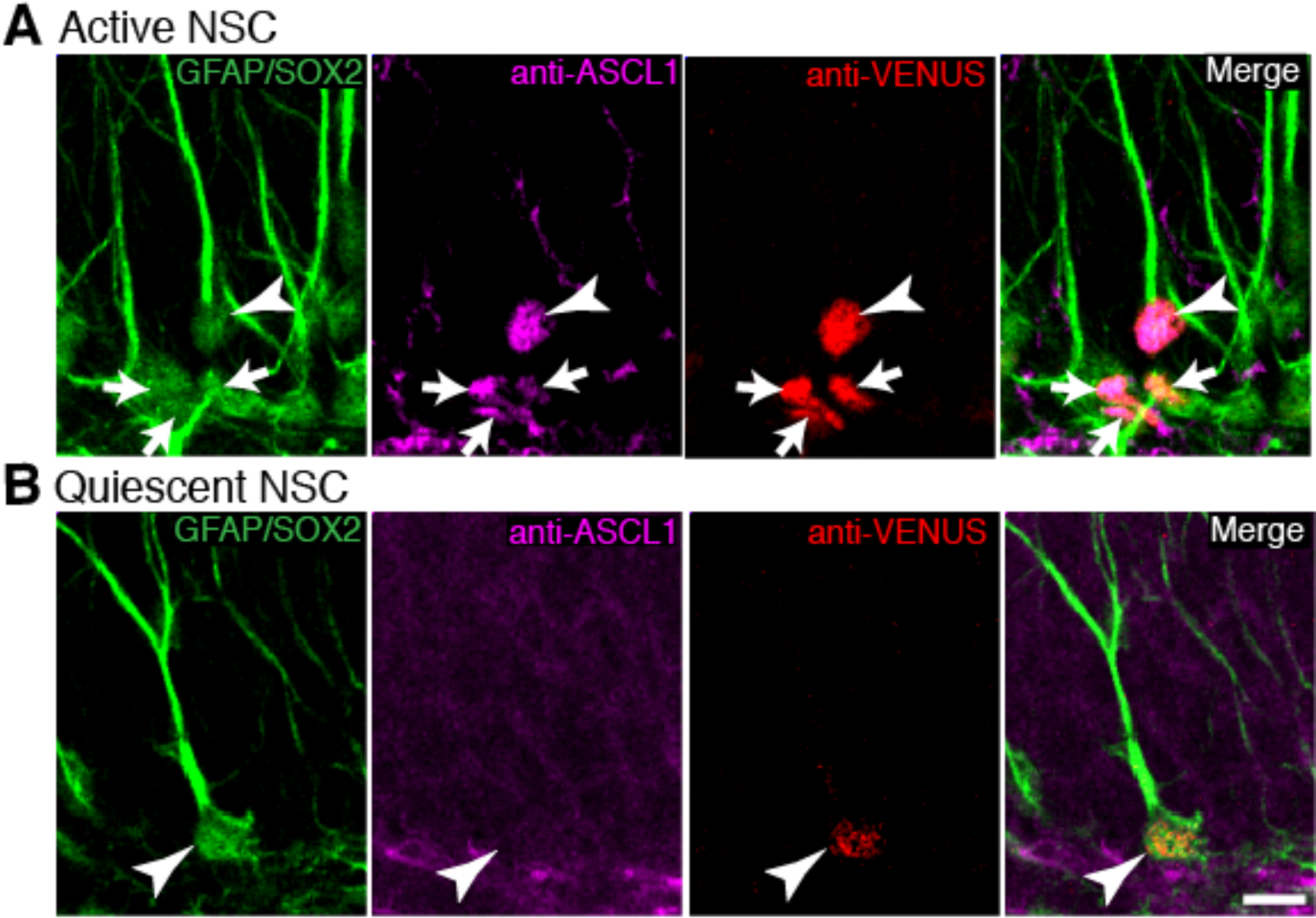
Increased detection sensitivity of ASCL1 protein using Ascl1-Venus mice and anti-GFP antibodies. (**A, B**) 3-week-old Ascl1-Venus mice, expressing a Ascl1-Venus fusion protein were stained for neural stem cell markers (GFAP and SOX2) as well as ASCL1 using both an anti-ASCL1 antibody and an anti-Venus/GFP antibody. (A) In active NSCs (active status indicated by penumbra of SOX2+GFAP-IPCs surrounding the NSC), ASCL1 protein was expressed at high levels and produced a strong signal using both antibodies. **(B)** In contrast, ASCL1 protein was expressed at very low levels in quiescent NSCs and could only be detected with the anti-Venus/GFP antibody. Scale bar (in **B**): **A**, **B** is 10 μm.

**Figure S7, related to Figure 7:**
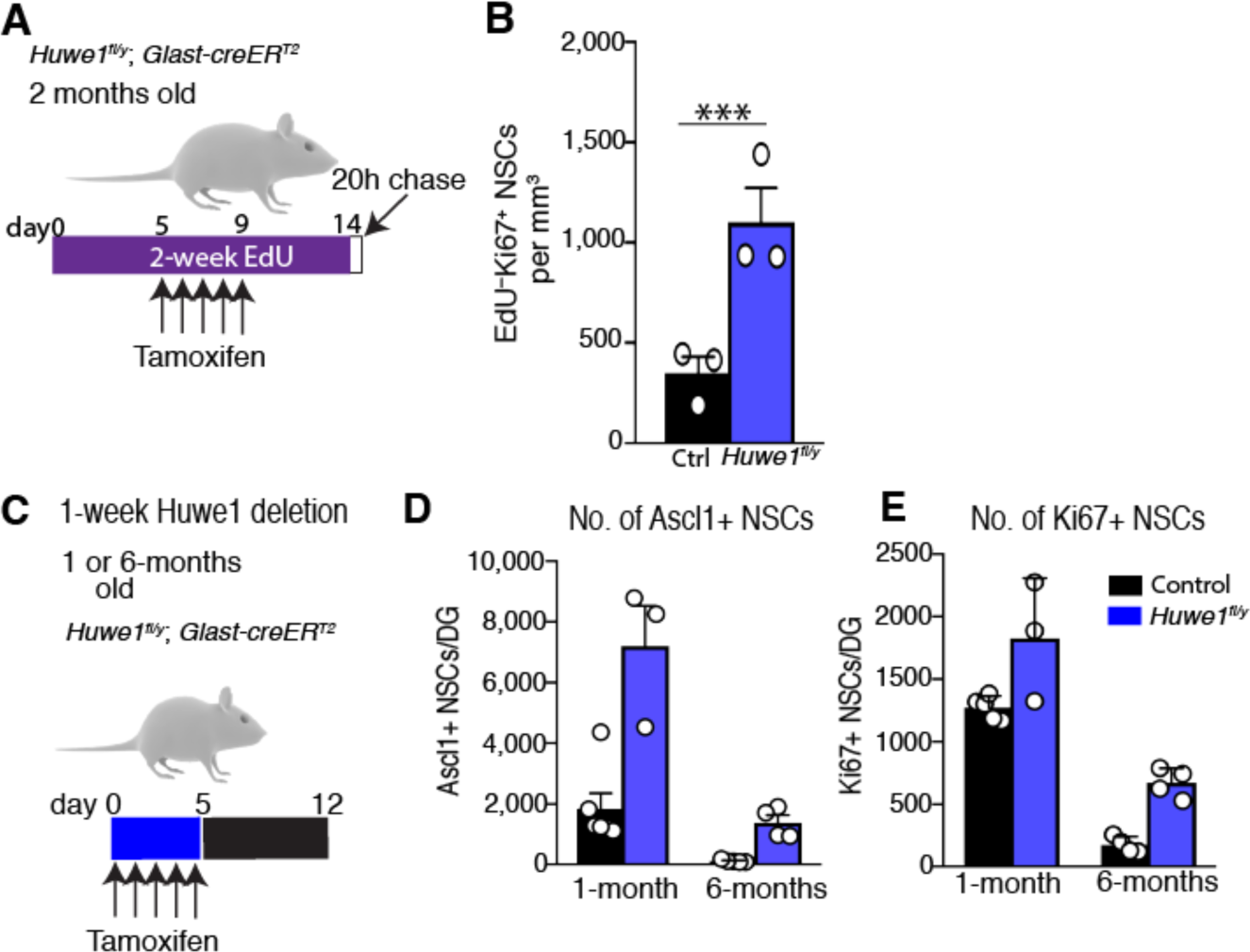
*Huwe1* suppresses the activation of dormant NSCs and increased activity of *Huwe1* with age (absolute counts). **(A**) *Huwe1^fl/y^*; *Glast-creER^T2^* mice (*Huwe1^fl/y^*) and controls were given EdU in drinking water for 2-weeks, during this time they also received tamoxifen. (**B**) Dormant NSCs (EdU-) in *Huwel’’* mice showed increased activation during the 16 h chase period (EdU-Ki67+) than in controls. (**C**) 1- and 6-month-old *Huwe1^fl/y^* and control mice were culled 1-week after receiving tamoxifen. (**D, E**) While the number of ASCL1+ and Ki67+ NSCs were increased upon *Huwe1* deletion at both ages, the effect was much larger in 6-month-old mice relative to age matched controls (see Figure 7 for graphs showing fold-change). Graphs represent mean ± s.e.m. Dots represent the mean of individual mice. Statistics: Student’s t-test. \*\*\**P*<0.001.

**Table S1, related to Figure 4:**
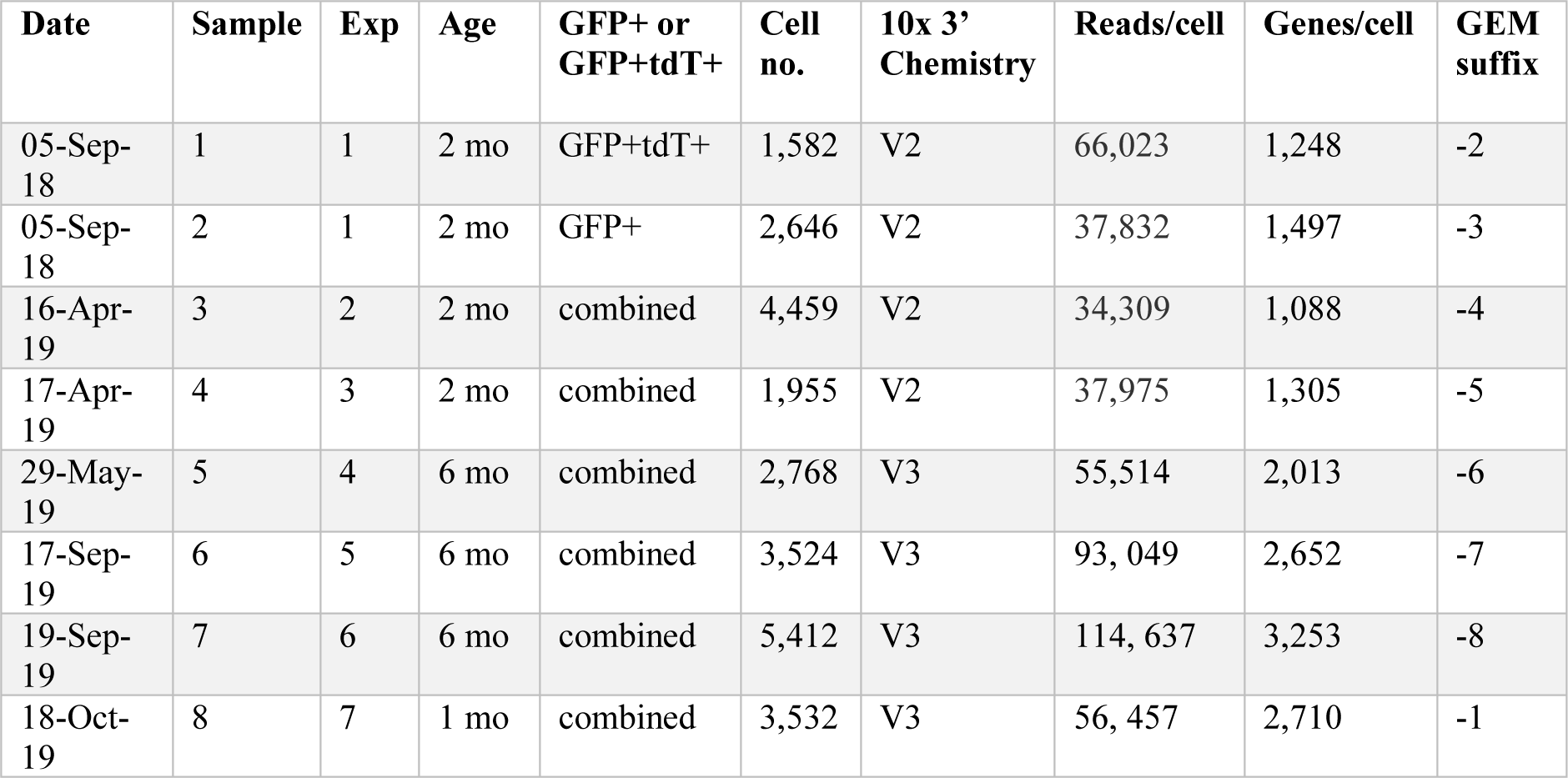
List of single-cell RNA sequencing experiments performed in Ki67-creERT2; lox-STOP-lox-tdTomato; Nestin-GFP mice.

| Date | Sample | Exp | Age | GFP+ or GFP+tdT+ | Cell no. | 10x 3' Chemistry | Reads/cell | Genes/cell | GEM suffix |
| --- | --- | --- | --- | --- | --- | --- | --- | --- | --- |
| 05-Sep-18 | 1 | 1 | 2 mo | GFP+tdT+ | 1,582 | V2 | 66,023 | 1,248 | -2 |
| 05-Sep-18 | 2 | 1 | 2 mo | GFP+ | 2,646 | V2 | 37,832 | 1,497 | -3 |
| 16-Apr-19 | 3 | 2 | 2 mo | combined | 4,459 | V2 | 34,309 | 1,088 | -4 |
| 17-Apr-19 | 4 | 3 | 2 mo | combined | 1,955 | V2 | 37,975 | 1,305 | -5 |
| 29-May-19 | 5 | 4 | 6 mo | combined | 2,768 | V3 | 55,514 | 2,013 | -6 |
| 17-Sep-19 | 6 | 5 | 6 mo | combined | 3,524 | V3 | 93,049 | 2,652 | -7 |
| 19-Sep-19 | 7 | 6 | 6 mo | combined | 5,412 | V3 | 114,637 | 3,253 | -8 |
| 18-Oct-19 | 8 | 7 | 1 mo | combined | 3,532 | V3 | 56,457 | 2,710 | -1 |

## 1 Model description

### 1.1 Purpose of the model

The aim of the model is to understand age-related changes of neural stem cell (NSC) numbers in the murine hippocampus. We distinguish between different NSC subpopulations, namely actively cycling NSC (identified by expression of Ki67) and quiescent NSC (identified by absence of Ki67 expression). The quiescent NSC population is further subdivided in dormant NSC, i.e., NSC that have never been activated since establishment of the niche at postnatal day 14 and resting NSC, i.e., quiescent NSC that have already been activated since establishment of the niche at postnatal day 14. The model describes the time evolution of active, dormant and resting NSC counts after postnatal day 14.

### 1.2 Data and experimental setup

Total, active and resting NSC counts were experimentally determined in mice aged 15, 35, 60, 200 and 365 days. For the measurements the animals were sacrificed such that each data point comes from a different individual.

NSC were counted in brain sections. Hippocampal NSC were defined as cells containing a radial GFAP-positive process linked to a SOX2-positive nucleus in the subgranular zone. Ki67 was used as a marker to distinguish between active (Ki67 positive) and quiescent (Ki67 negative) stem cells.

The resting NSC counts were approximated by the following experimental procedure: Mice were exposed to the thymidine analogue EdU for 14 days followed by an EdU free period of 20 hours (referred to as ”chase”). During DNA repliction EdU is incorporated into the DNA. After the chase period the animals were sacrificed. Resting cells were defined as EdU positive Ki67 negative cells, i.e., cells that have divided during the EdU exposure but have returned to a quiescent state at the time of cell counting.

### 1.3 Model derivation

The model describes time evolution of active, resting and dormant NSC. As stated above, NSC that have never divided during post-embryonic life are denoted as dormant. Quiescent NSC that have divided during post-embryonic life are denoted as resting NSC. Cycling stem cells are denoted as active. The model considers the following processes

- Dormant NSC are activated at the rate r_1_.
- Resting NSC activated at the rate r_2_.
- Active stem cells divide at the rate p.
- Upon division an active NSC gives rise to two progeny. With probability a a progeny cell is again a stem cell (referred to as self-renewal), with probability (1 - a)itisan intermediate neuronal progenitor cell (referred to as differentiation). The probability a is referred to as self-renewal probability or fraction of self-renewal [3, 5, 6].
- In agreement with our previous work and with experimental data [2] we assume that NSC originating from division become quiescent.

The model is summarized in Figure 1. It is an extension of previous models [2, 7, 8]. The new aspect of the model introduced here is the distinction between resting and dormant NSC.

**Figure 1:**
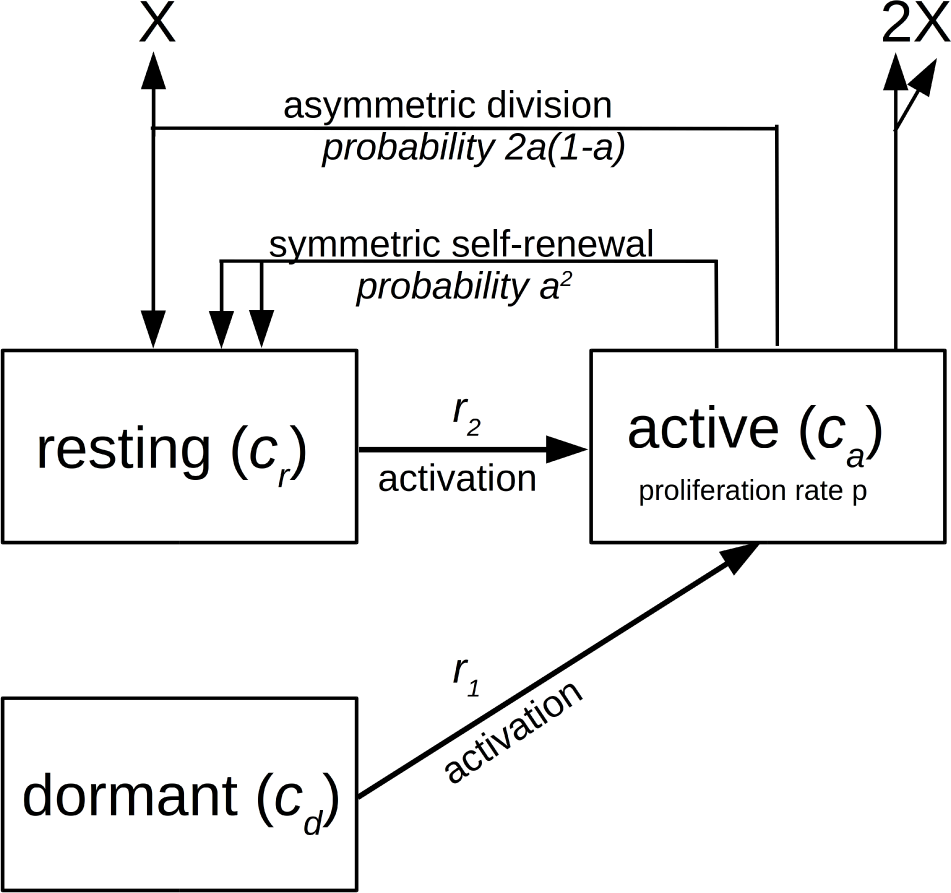
NSC model. The scheme shows processes described by system (1). Non-stem cells are denoted by X

The rate r_2_ describes the reactivation from the resting state. If r_2_ assumes very large values this corresponds to a scenario where the time spent in the resting state before reactivation is negligibly short. In biological terms this means that most cells remain active after division.

We denote the amount of dormant stem cell at time t as c_d_(t). The amount of active stem cells at time t is denoted as c_a_(t) and that of resting stem cells as c_r_(t). For notational convenience we omit the argument t and identify *c_d_(t) = c_d_, c_r_*(t) = c_r_ and c_a_(t) = c_a_. This results in the following system of ordinary differential equations.

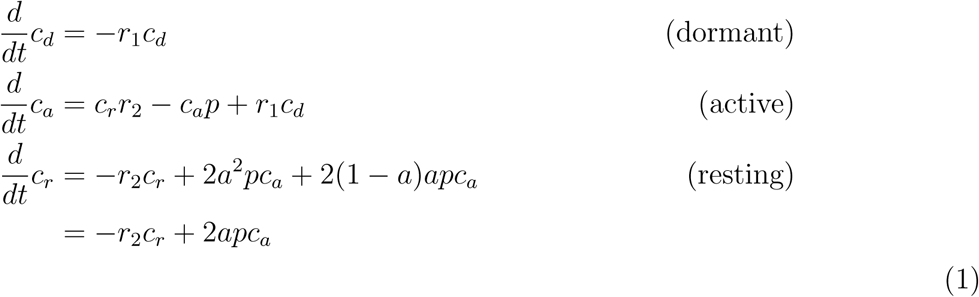

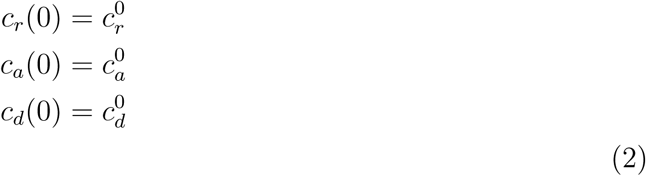

with nonnegative initial conditions c_r_^0^, c^0^_d_ and c^0^_a_.

We note that for r_1_ = r_2_ and qNSC = c_d_ + c_r_, where qNSC denotes the amount of quiescent stem cells, we obtain the model from [2].

### 1.4 Model of the labeling experiments

Experimentally we can only count the resting cells that have divided during the EdU exposure and became quiescent afterwards. The obtained cell counts are a lower bound for the total number of resting cells, since resting cells that have not divided during EdU exposure cannot be identified in the experiment. To avoid potential underestimations we explicitly simulate the experimental setup and compare the experimentally obtained resting cell counts to the simulations.

#### 1.4.1 Labeling phase

For the labeling phase we assume:

- During EdU supply all active cells are instantaneously labeled.
- Cells transiting during the time of EdU supply from active to resting state retain the label.

To quantitatively describe this experiment we have to distinguish between labeled and unlabeled resting cells. Since we assume that active cells get instantaneously labeled all active cells are per definition labeled. During the EdU supply the following model is considered. It is visualized in Figure 2. We denote as C_d_ the amount of dormant unlabeled cells, as *C_r,labeled_* the amount of resting labeled cells, as *C_r,unlabeled_* the amount of resting unlabeled cells and as C_d_ the amount of active labeled cells.

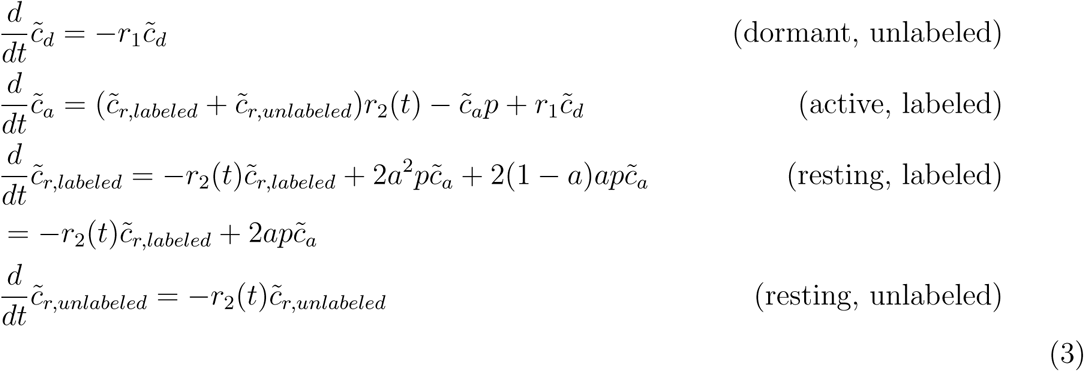

**Figure 2:**
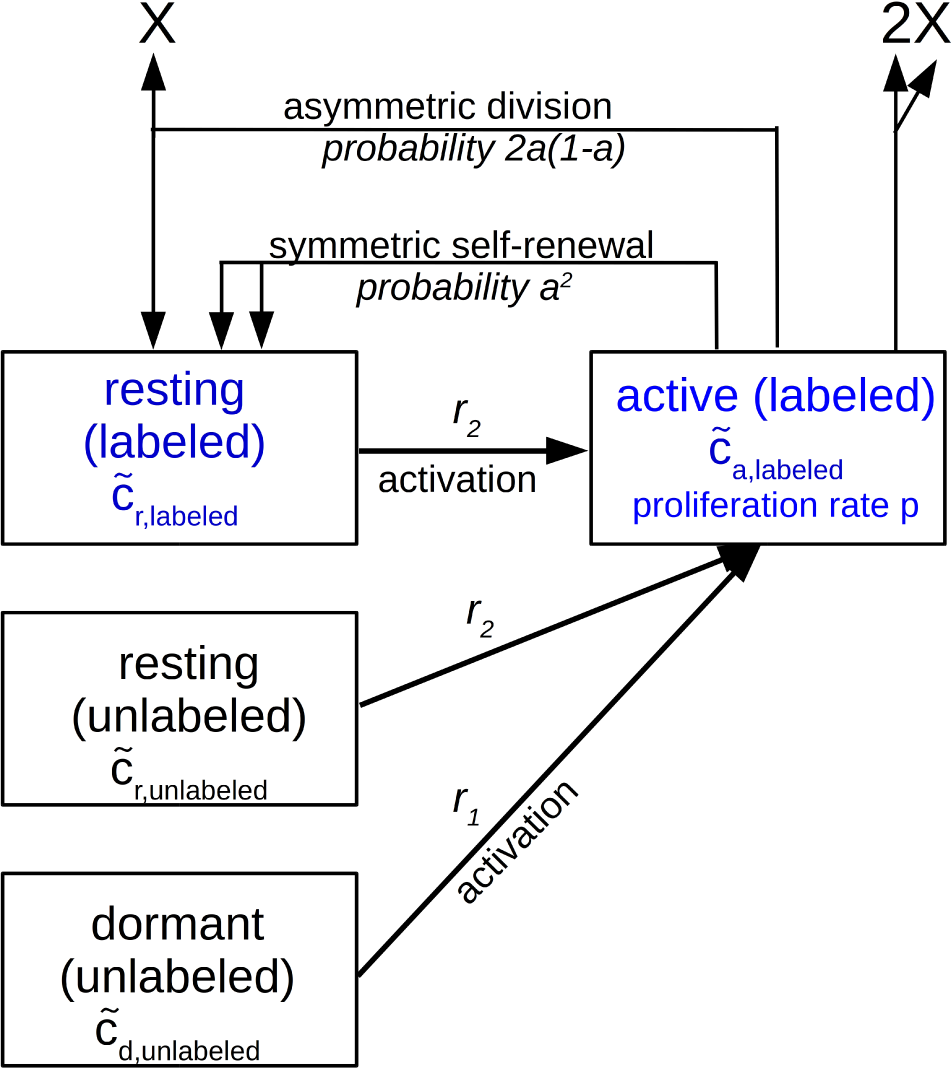
NSC model during EdU supply. The scheme shows processes described by system (3). Non-stem cells are denoted by X.

Assume the EdU supply starts at *t = t*,* then we have

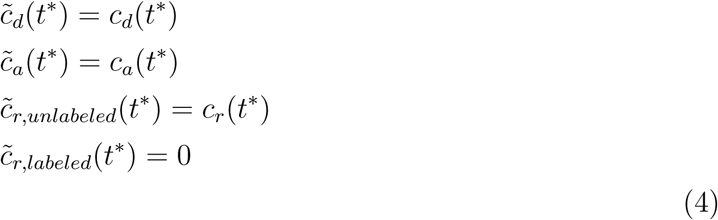

#### 1.4.2 Chase period

For the chase period we make the following assumptions.

- Unlabeled cells getting activated during the chase period remain unlabeled and produce unlabeled offspring.
- Since the chase period is short and labels are retained during several divisions, we assume that labeled cells dividing during the chase period give rise to labeled cells.
 This leads to the following system of equations. We denote as *C_r_,_labeled_* the amount of resting labeled cells and as *Ca,labeled* the amount of active labeled cells. The model is visualized in Figure 3. 

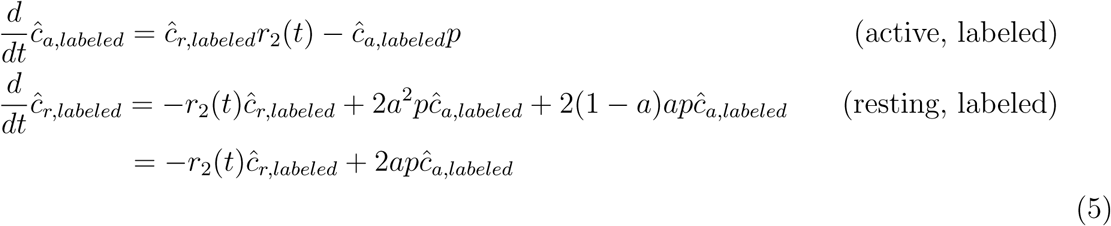

**Figure 3:**
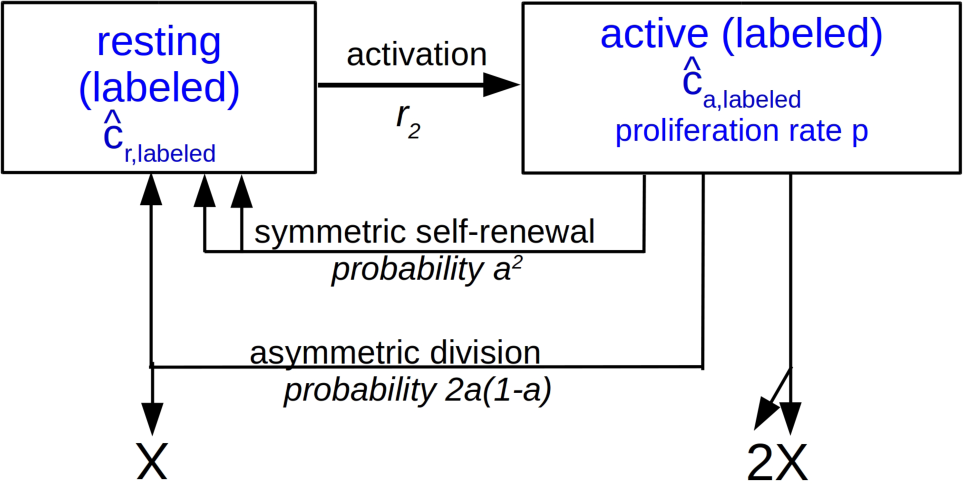
NSC model during the chase period. The scheme shows processes described by system (5). Non-stem cells are denoted by X.

Assume the chase starts at t — t^#^,thenwehave 

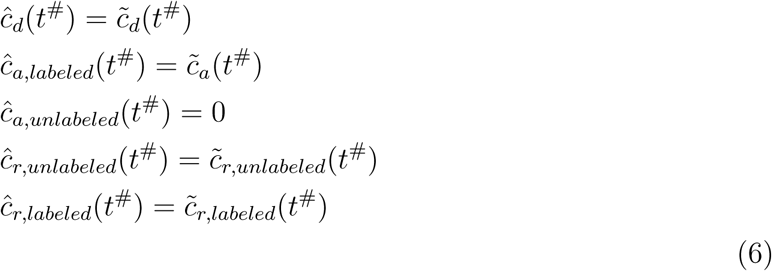

### 1.5 Full model

The amount of resting (EdU+Ki67-) cells present at time t is obtained as follows. Time evolution of active, resting and dormant NSC before EdU tem (1) with initial conditions (2). We start simulation at time 0 and stop at *t* — t.* Then we simulate the labeling period of *T_EdU_* — 14 days using system (3) with the initial condition (4) for *t*\* — *t*. We then simulate the chase using system (5) with the initial condition (6) for t# — t* + *T_EdU_*. We stop simulations at t — t* + *T_EdU_* + *T_chase_,* with *T_chase_* — 20 hours to readout the resting labeled cell count to compare it to measurements. The procedure is summarized in Figure 4.

**Figure 4:**
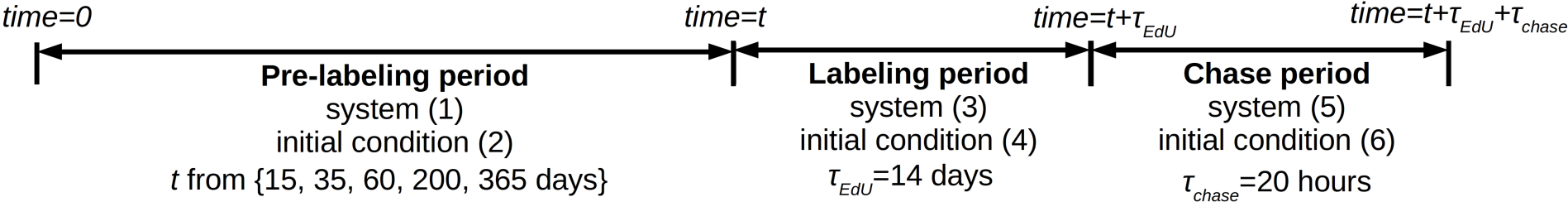
Simulation of labeling experiments.

## 2 Model simulations

The model has been implemented in MATLAB and ordinary differential equations have been solved using the solver ode23s.

## 3 Model quantification

As in [2] we assume a doubling time of active NSC of 22.8 hours, as measured in [4]. The other model parameters and the initial conditions c^0^_a_, c_r_^0^, c^0^_d_ are fitted based on the data. For fitting we use a multistart approach (5000 multistarts) with random nonnegative initial parameter guesses. The sampling of the initial guesses follows a latin hypercube approach. Optimization is performed using the MATLAB function fmincon.

## 4 Model selection

To compare different models we use the Akaike information criterion for small sample sizes (AI Cc) given as 

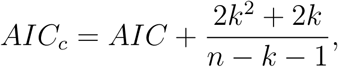

where n is the number of data points, k the number of free parameters and AI C the Akaike information criterion [1]. This approach takes into account the accuracy of the fit but at the same time punishes a high number of free parameters to prevent overfitting. The level of empirical support of a given model is considered substantial if 0 < A < 2, considerably less so if 4 < A < 7 and none, if A > 10 holds, where A is the AIC_c_ of a given model minus the minimum AICc of all considered models [1].

## 5 Age-related change of cell parameters

In [2] it has been concluded that the activation rate of NSC changes during aging. This can be modeled by assuming that the rate is not given by a constant but by a time-dependent function. In the following we allow the activation rates r1, r2 and the self-renewal probability a to be functions of the age of the organism.

## 6 Fitting active and total NSC dynamics

To check consistency of our dataset with the model from [2] we first fit different versions of the model only to active and total NSC counts, i.e., for the moment we ignore the information about resting NSC.

### 6.1 Identical activation rates for dormant and resting NSC

In [2] it has been concluded that the activation rate of quiescent NSC changes during aging. To obtain the model from [2] we assume that the activation rates are identical for dormant and resting cells and that they change with age. As in [2] we set 

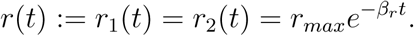

As in [2] we assume a cell cycle time of active NSC of 22.8 hours, as measured in [4]. We fit the other model parameters to active and total NSC counts. We observe that the model is close to the data, however, the values of the estimated parameters differ from those in [2].

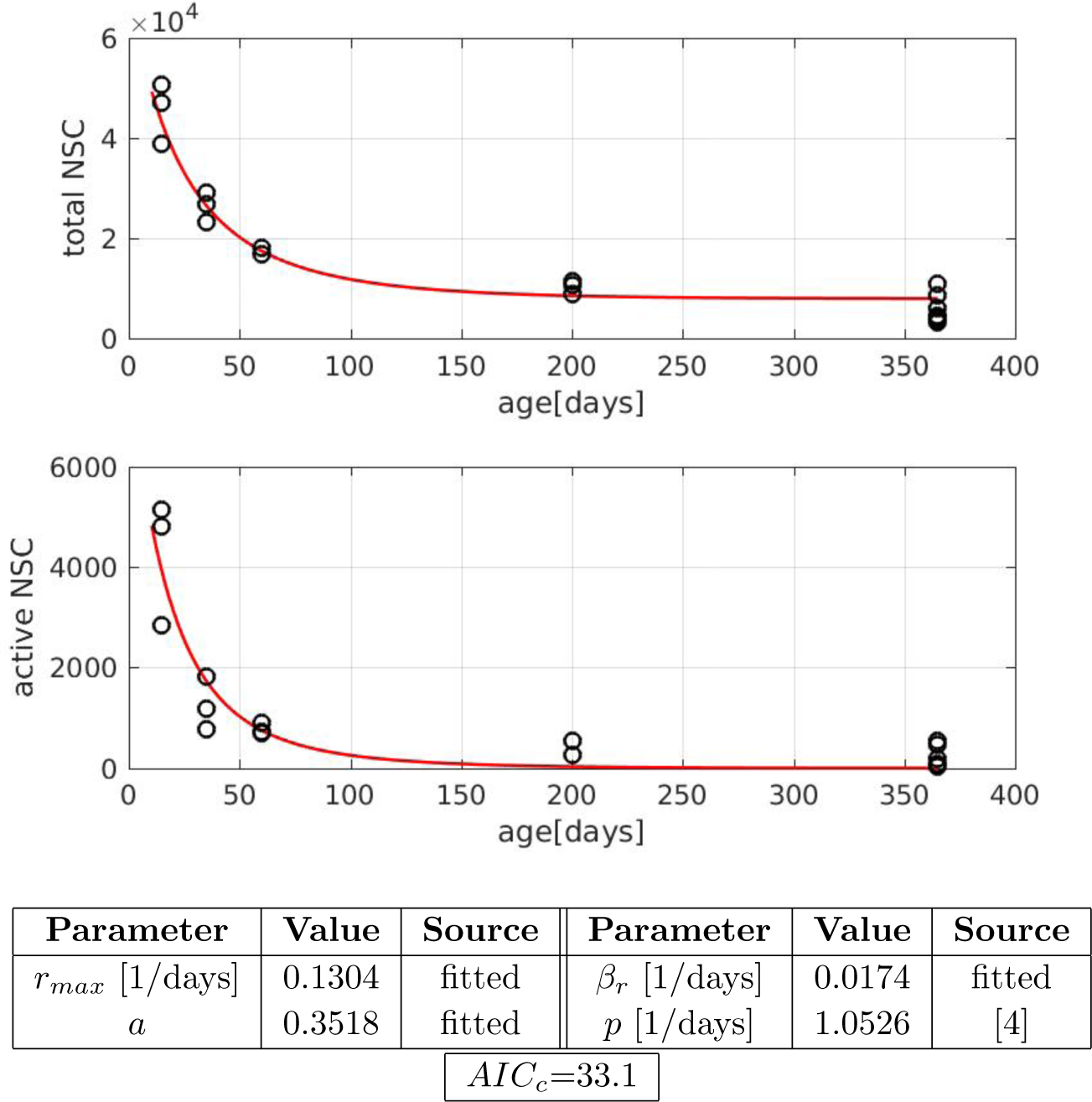

### 6.2 Different activation rates for dormant and resting NSC

We now consider the case where activation rates of dormant and resting NSC can be different. As in [2] we assume that they decline exponentially with time, i.e., 

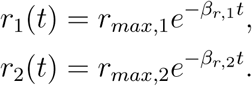

We fit this version of the model to the counts of total and active NSC. This does not improve the fit and an increase in AICc demonstrates that based on the total and active cell counts there is no evidence for the rates to be different.

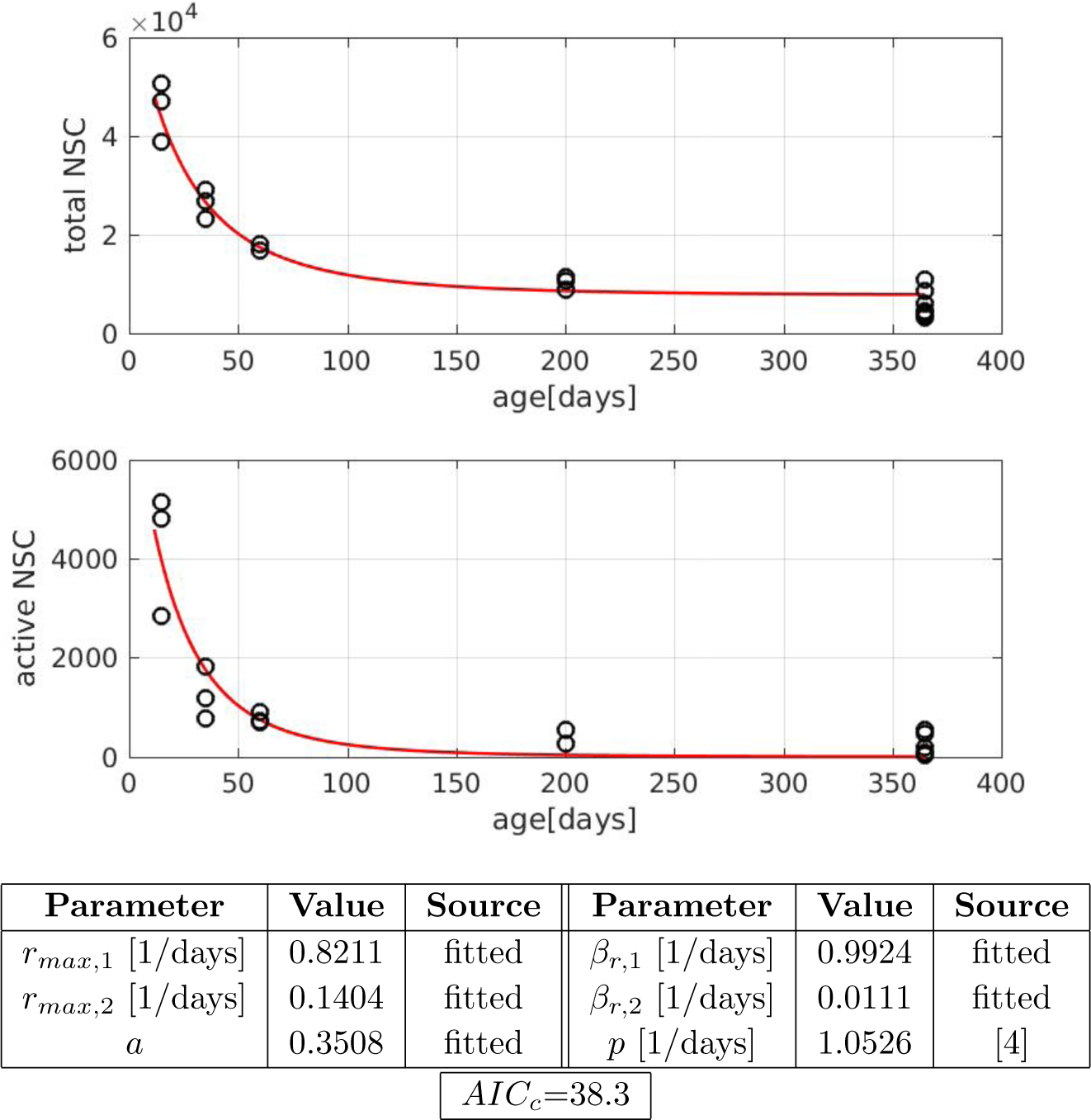

## 7 Fitting total, active and resting NSC dynamics

Now we include our data on resting NSC. In this section we fit the model to total, active and resting NSC counts.

### 7.1 Identical activation rate for dormant and resting NSC

We first consider the scenario where dormant and resting NSC have the same activation rate, i.e., we set 

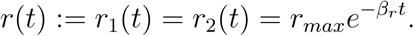

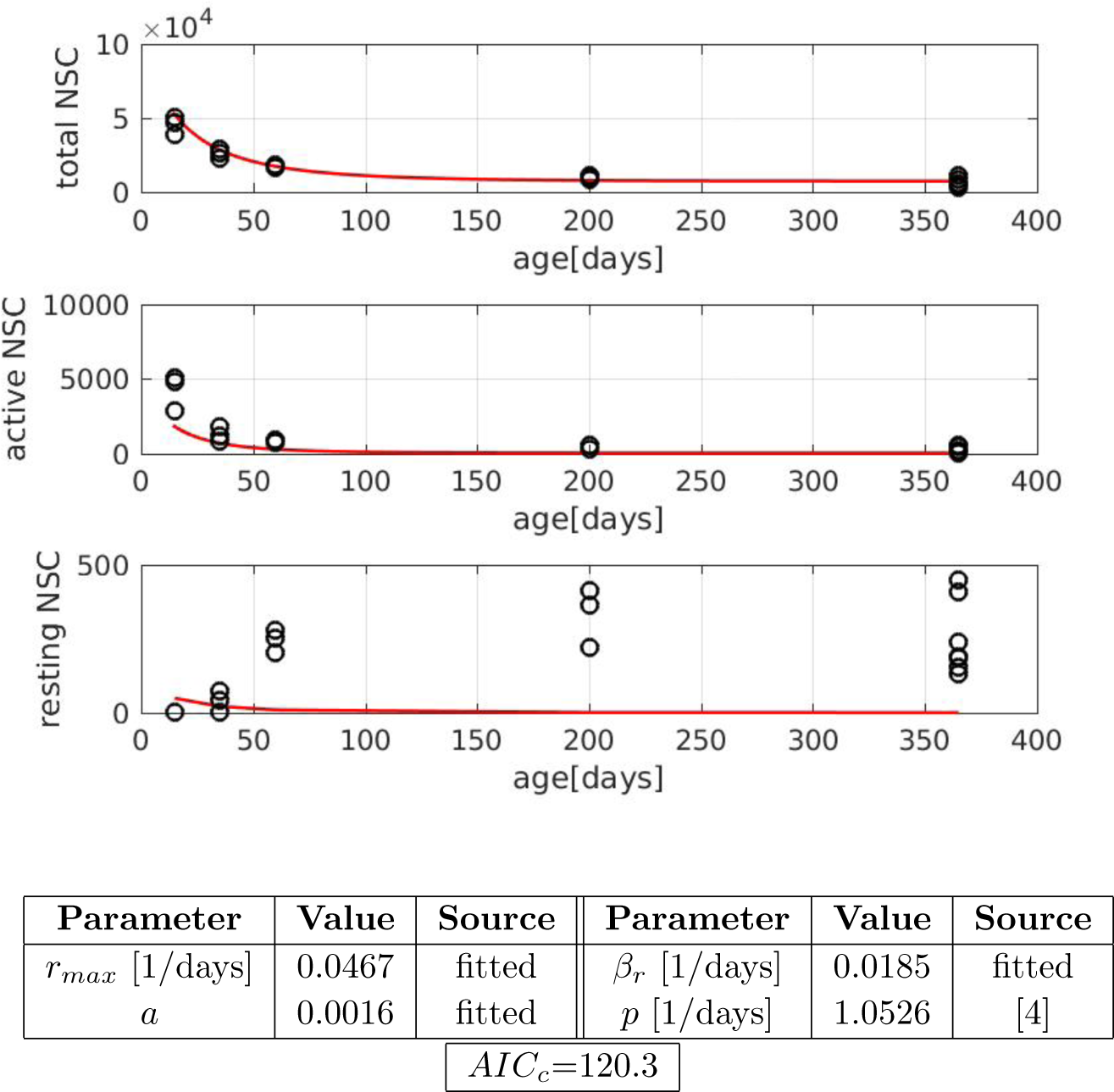

### 7.2 Different activation rates for dormant and resting NSC

If we allow dormant and resting cells to have different activation rates, i.e., 

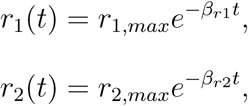

we obtain a better fit and a reduction in AICc. This suggests that different activation rates are required to explain the observed dynamics of resting NSC.

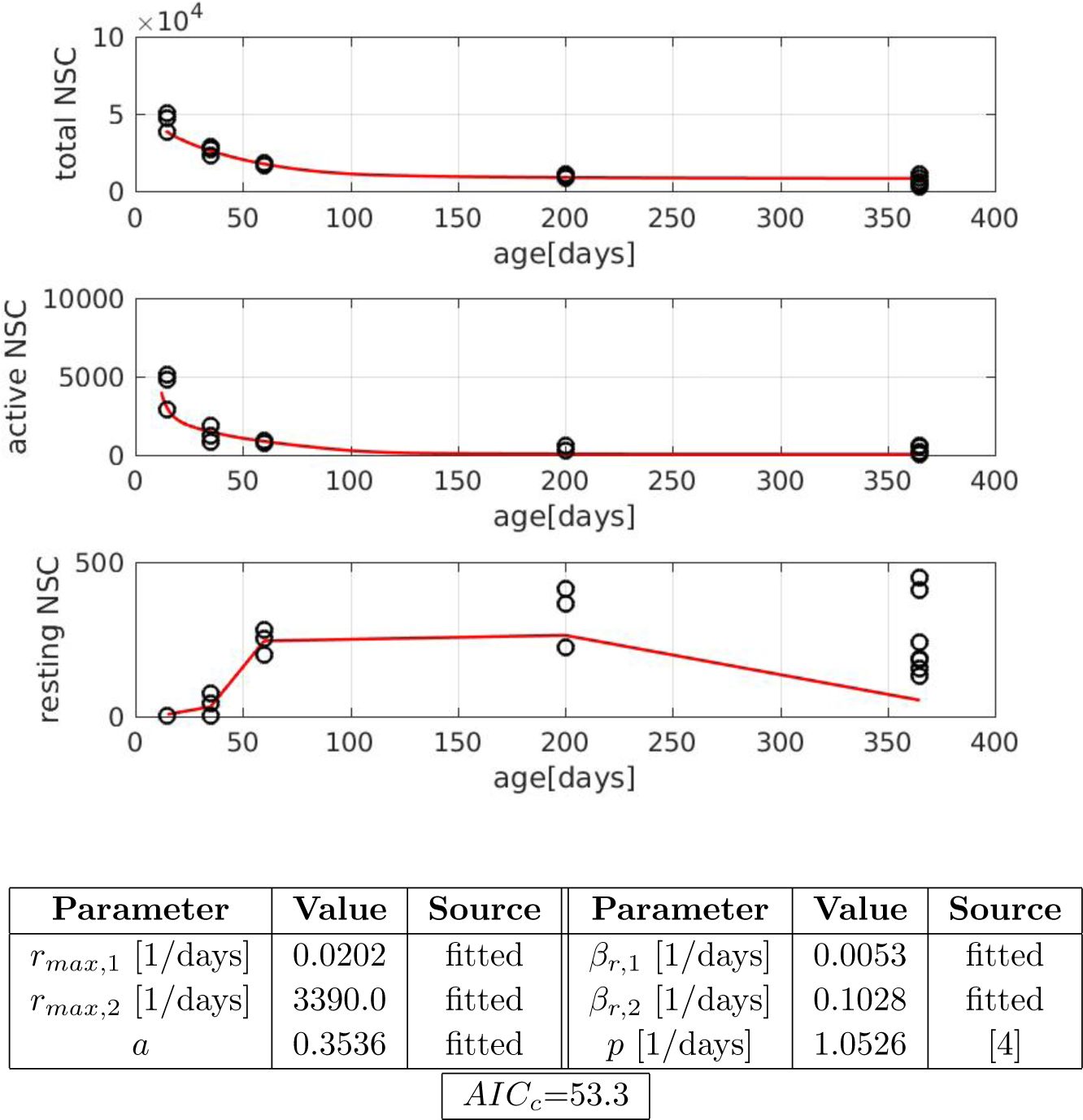

The high activation rate of resting cells indicates that in young mice most activated cells do not return to quiescence after division.

### 7.3 Different activation rates for dormant and resting NSC and age dependent self-renewal

We now allow different activation rates for dormant and resting cells and we allow self-renewal to increase over time. The expression for age-dependent self-renewal has been taken from [2]. We set 

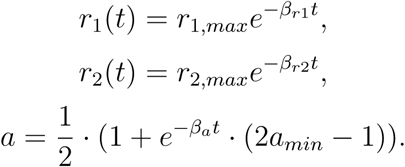

This best obtained fit is the following.

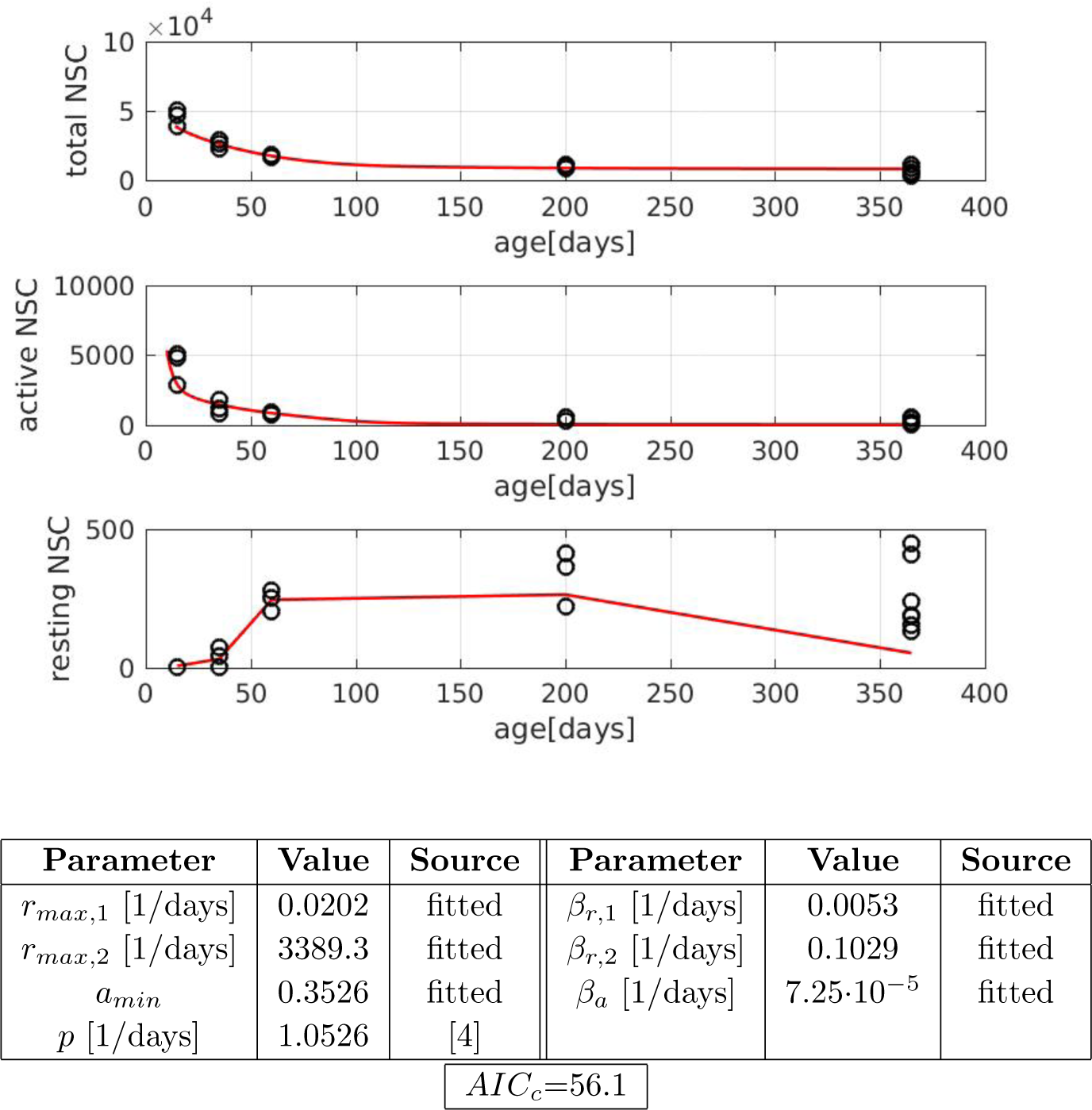

We observe an increase in AICc supporting the view that age dependence of self-renewal has only a minor impact on cell dynamics. Also here the high activation rate of resting cells indicates that in young mice most activated cells do not return to quiescence after division.

The following figure shows how activation rates and self-renewal probability change with age.

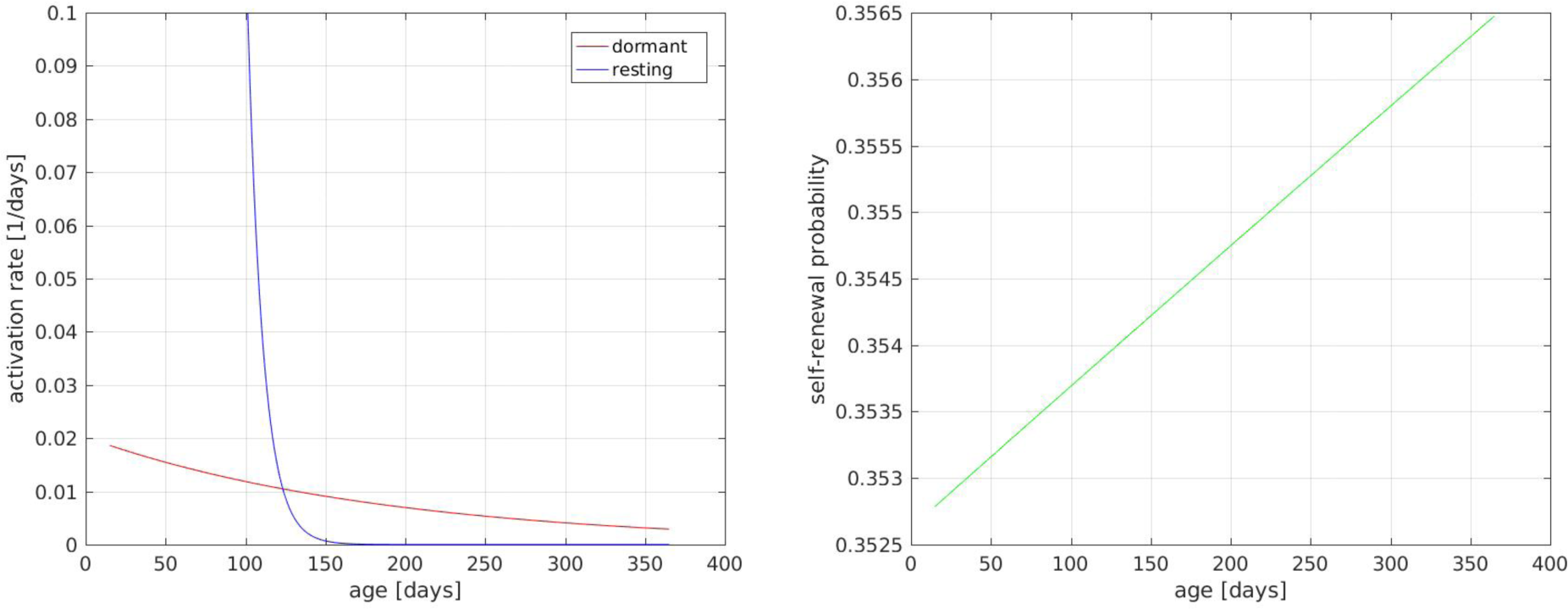

### 7.4 Model with non-zero activation for large time

In the versions of the model considered so far, the activation rates decline asymptotically with time. This can lead to rates that are practically zero within the life-time of a mouse. To prevent this unrealistic scenario, we modified the model as follows: 

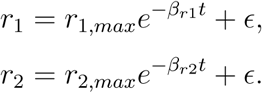

This implies that the activation rate is always larger than e.Wefite in addition to the other parameters.

#### 7.4.1 Constant self-renewal

We first consider the scenario with constant in time self-renewal (a = const) and age-dependent activation rates: 

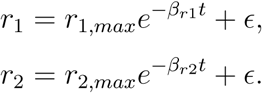

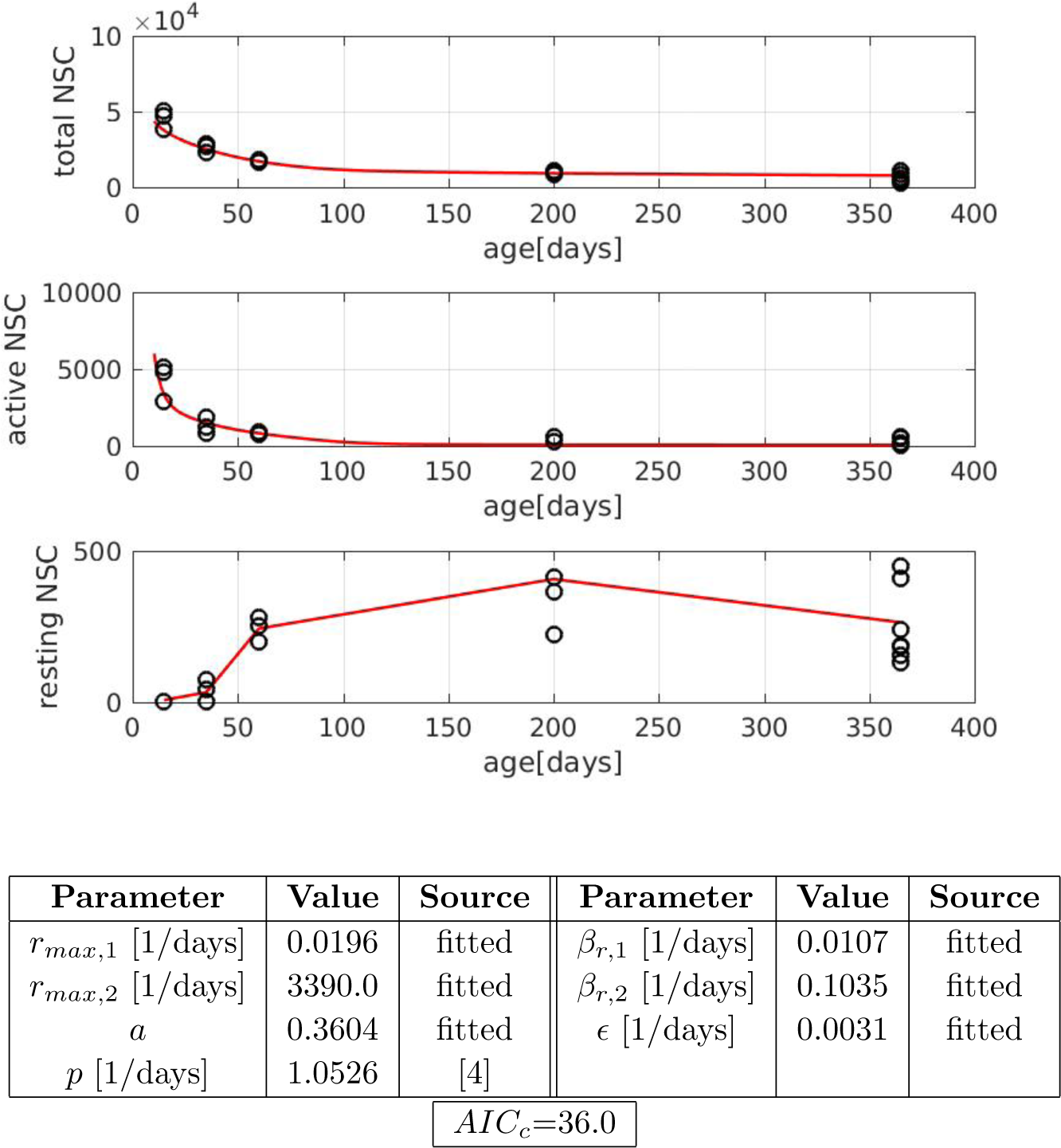

The fit is significantly improved. Although we have one more free parameter AI C_c_ is reduced compared to the previous versions of the model. This implies that it is important for the observed process that activation rates do not decline to zero for high ages.

#### 7.4.2 Age-dependent self-renewal

We now allow in addition that self-renewal is age-dependent. As motivated above, we set 

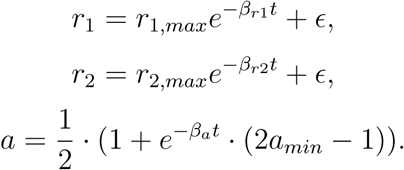

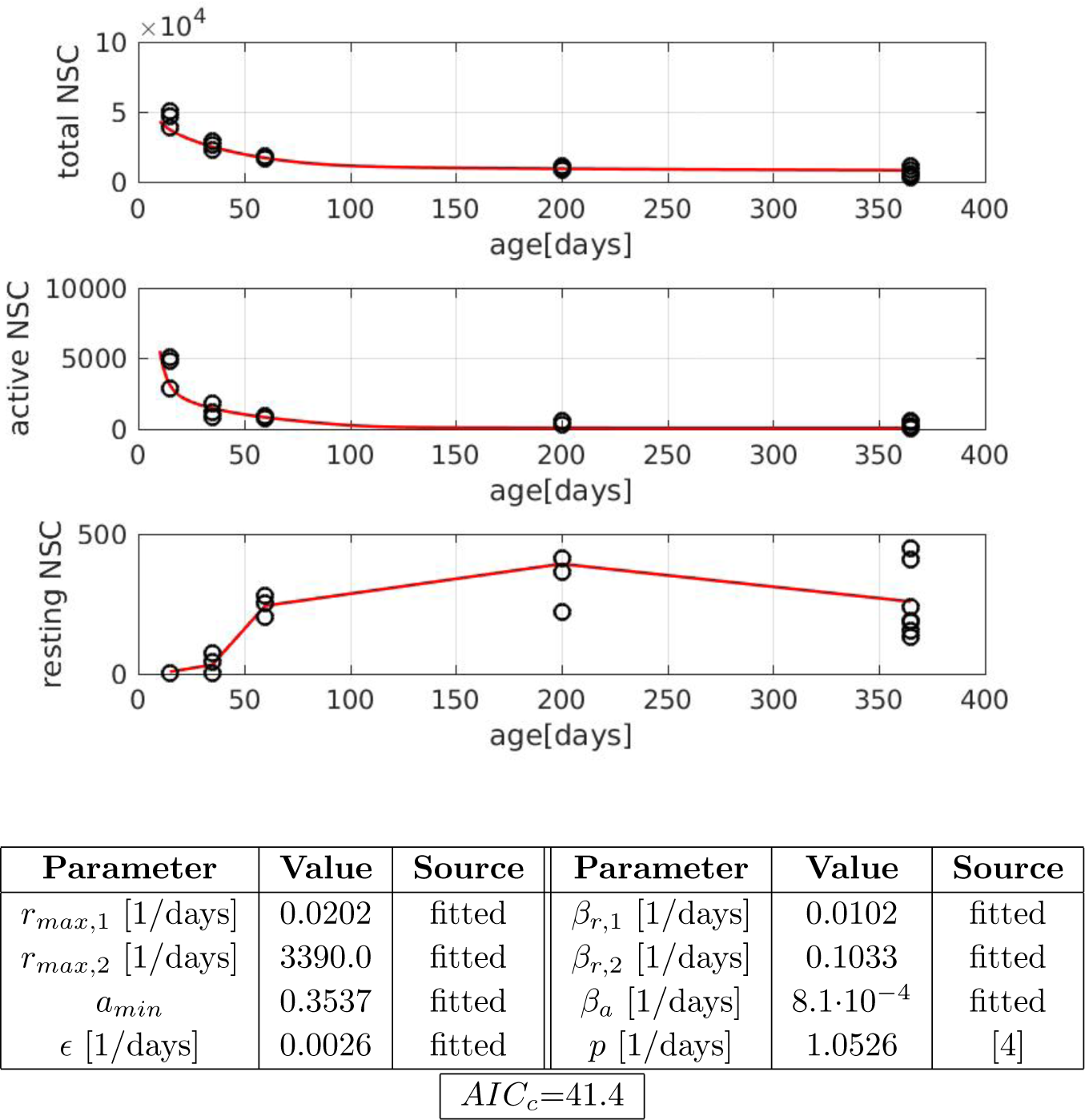

The increase in AI Cc indicates that age-dependence of self-renewal has only little impact on the NSC dynamics.

The following figure shows how activation rates and self-renewal probability change with age.

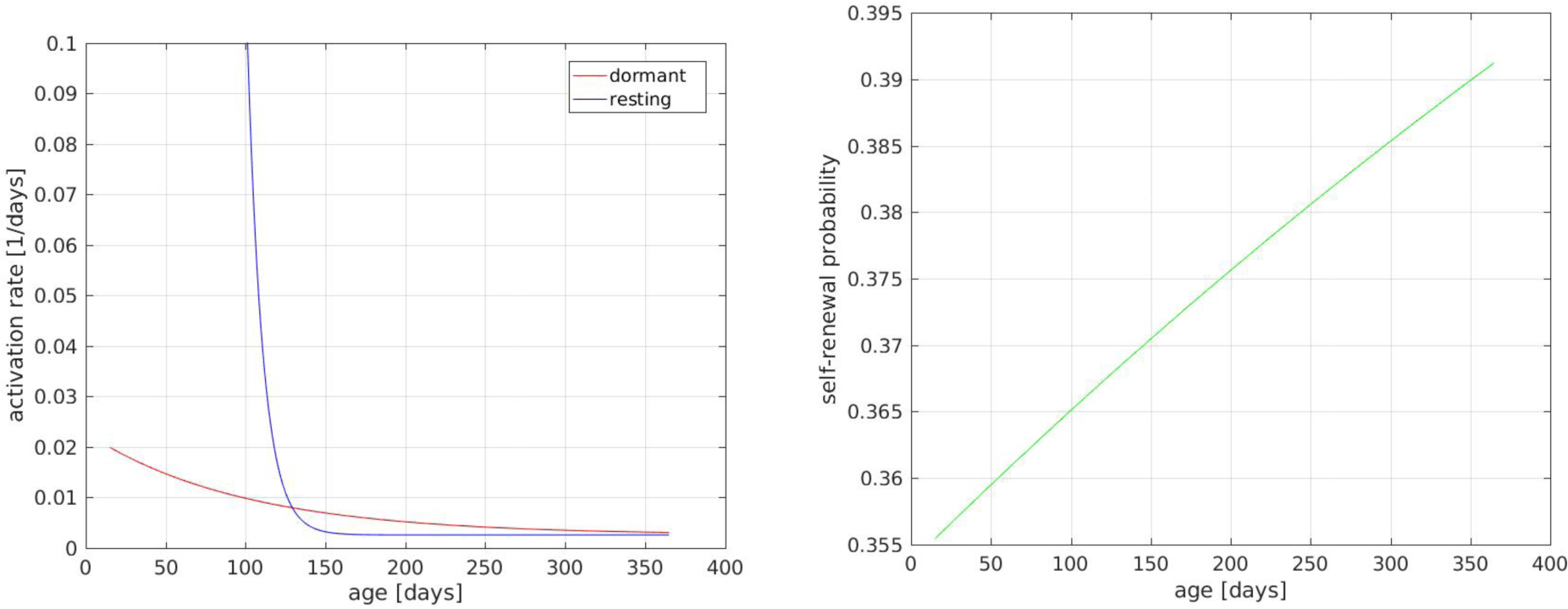

